# Large scale phage-antibiotic combination studies reveal key combinations for urinary tract infection and urosepsis treatments

**DOI:** 10.64898/2026.03.10.709588

**Authors:** Karen D. Adler, Slawomir Michniewski, Caitlin Wildsmith, Eleanor Jameson, Nathan Brown, Aimee M. Daum, Mahmuda Akter, Marie Attwood, Jennifer Mahony, Ozcan Gazioglu, J. Mark Sutton, Martin Textor, Thomas Sicheritz-Pontén, Andrew D. Millard, Melissa E. K. Haines, Martha R J Clokie

## Abstract

The growing problem of AMR infections in healthcare has prompted the search for alternative treatments, with increasing interest in bacteriophages. However, most bacteriophage-antibiotic interactions are incompletely understood, and the benefits of combining them remains context dependent.

In this study, we screened thousands of phage-antibiotic combinations to assess interaction outcomes in clinical *E. coli* and *K. pneumoniae* isolates. By integrating bacteriophages into an existing, scalable clinical MIC determination platform, we identified shifts in antibiotic MIC and susceptibility, revealing patterns of additivity and antagonism.

Overall, interactions showed a species-specific profile; additive interactions predominated, particularly for *E. coli*. Hierarchical clustering highlighted frequent positive interactions between β-lactams and Tequatroviruses. Notably, closely related phages sometimes displayed divergent phenotypes, indicating that interaction outcomes cannot be inferred solely from taxonomic relatedness or genomic similarity. Taken together, these results establish a foundation for rational, evidence-based development of phage-antibiotic therapies to restore and broaden treatment options against multidrug-resistant infections.

## Introduction

Antimicrobial resistance (AMR) is a major and escalating threat to global health. In 2019, 1.27 million deaths were directly attributable to AMR, with a further 4.95 million deaths associated with resistant infections^1^. Despite ongoing efforts to develop new antimicrobials, little progress has been made, and treatment options for many bacterial infections continue to diminish, increasingly forcing reliance on older, more toxic antibiotics such as colistin, even as resistance to these last-line agents emerges^2^.

Urinary tract infections (UTIs) are among the most common bacterial infections worldwide and are increasingly associated with antibiotic resistance. Between 50-80% of women experience at least one UTI during their lifetime, and up to half develop recurrent infection^3^. UTIs are also a major source of complications, accounting for approximately 20% of hospital-acquired bloodstream infections (BSIs), with a mortality rate of ∼10%^4^. The predominant causative agents, *Escherichia coli* and *Klebsiella pneumoniae*, are both designated Critical Priority pathogens by the World Health Organization. In the EU/EEA, resistance was detected in 54% of invasive *E. coli* isolates and 38% of *K. pneumoniae* isolates in 2022, with multidrug resistance observed in 10.8% and 22.1% of isolates, respectively^5,6^.

Pathogenic *E. coli* display a highly conserved population structure dominated by a small number of globally disseminated sequence types with clade B2 (ST131, ST73, ST95, ST127) and clade D (ST69) dominating UTIs, urosepsis, and BSIs across diverse settings^7–17^. In contrast, pathogenic *Klebsiella* spp. show no comparable clonal dominance, reflecting fundamental differences in pathogen ecology and evolution^18^. These features make *E. coli* and *K. pneumoniae* complementary models to study antimicrobial intervention strategies.

As resistance increases, UTIs that were once readily managed with short oral antibiotic courses now frequently require hospitalisation and prolonged intravenous therapy, with a corresponding rise in difficult-to-treat BSIs. Strategies that result in bacterial eradication or prevent persistence, recurrence, or progression of these infections are therefore of increasing clinical importance.

Bacteriophage (phage) therapy has re-emerged as a potential adjunct to antibiotic treatment, owing to its specificity for bacterial hosts, activity against multidrug-resistant strains, capacity to disrupt biofilms, and favourable safety profile^19^. However, antibiotics remain the only widely established and regulated antimicrobial therapy in clinical use. Combining phages with antibiotics therefore represents a rational approach to leverage complementary antibacterial mechanisms rather than replacing existing treatments.

Previous studies have reported both synergistic^20–25^ and antagonistic^26–30^ phage-antibiotic interactions, introducing the phenomenon of phage-antibiotic synergy (PAS). However, most investigations have examined a small number of phages and antibiotics, often under heterogeneous experimental conditions, restricting the ability to identify generalisable interaction principles. In addition, reliance on case reports and small-scale studies has further constrained mechanistic and translational insight due to specific nuances related to phage and antibiotic selection, as well as lack of dosing rationale for the combination of phages and antimicrobial concentrations, frequency, and duration of therapy.

Here, we systematically investigate phage-antibiotic interactions at scale by screening thousands of combinations across large phage collections and twenty-four antibiotics, using multiple patient-derived, highly resistant *E. coli* and *K. pneumoniae* isolates. By embedding phages within a clinically relevant antimicrobial susceptibility testing framework, we define interaction outcomes and identify species-specific patterns that inform the rational development of combination therapies.

## Results

### Bacteria and Phage Panel Refinement

An overview of the experimental workflow and outcomes is shown in **Figure 1**.

**Figure 1:**
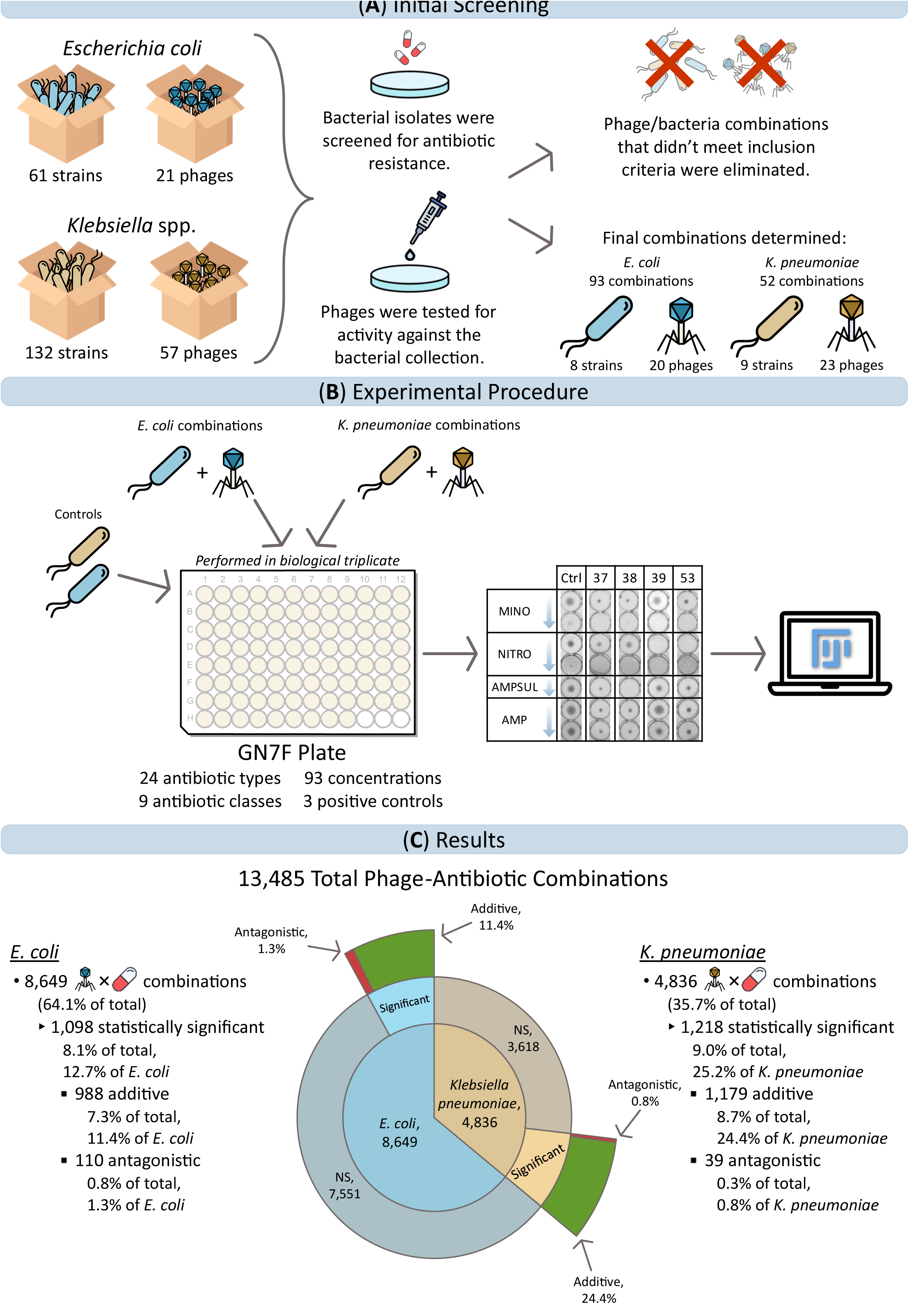
Experimental workflow and interaction outcomes. (A) Clinical *E. coli* and *Klebsiella* spp. isolates from UTIs, urosepsis, and BSIs were assembled, alongside a large phage panel targeting each species. Phages unable to achieve high titre propagation (10^8^ PFU·ml^-1^) or demonstrate infectivity were excluded as were bacterial isolates intrinsically antibiotic-sensitive or phage resistant. (B) Bacterial isolates were inoculated into Sensititre GN7F plates (antibiotic-only controls). Parallel plates were prepared with phage added at a multiplicity of infection (MOI) of 1. Assays were performed in biological triplicate. Bacterial biomass was quantified using ImageJ and phage-antibiotic combinations were compared to matched antibiotic-only controls. (C) Across 13,485 measurements, 2,316 interactions (17%) showed significant deviation from control. Most were additive (2,167; 16%), with relatively few antagonistic interactions detected (149; 1%).

To assess how bacteriophage-antibiotic combinations influence antibiotic efficacy, we first identified bacterial isolates that were resistant to a broad range of antibiotics yet susceptible to multiple phages.

A total of 61 *E. coli* and 132 *Klebsiella* spp. clinical isolates (**Supplementary Tables 1** and **2**) were screened against 21 coliphages and 57 *Klebsiella* phages, respectively (**Supplementary Tables 3** and **4**).

Bacterial isolates were excluded if they showed low or no phage efficacy (26 *E. coli* [∼43%]; 104 *Klebsiella* spp. [∼79%]), or were sensitive to multiple antibiotics (26 [∼43%] *E. coli* ; 15 [∼11%] *Klebsiella* spp.). From the remaining 22 strains, 17 were selected: eight *E. coli* and nine *K. pneumoniae* strains **(Supplementary Table 5**).

The remaining *E. coli* isolates spanned three clades (B1, B2, and D) and five STs (38, 73, 131, 357, and 940), all associated with human infection (**Figure 2A**), including the globally disseminated MDR lineage ST131^31^. The remaining *K. pneumoniae* isolates belonged to phylotype Kp1, (*K. pneumoniae* sensu stricto) and five ST types (11, 147, 258, 307, and 661) (**Figure 2B**). Genomic interrogation revealed strong concordance between predicted AMR determinants and phenotypic resistance, with all strains resistant to 7-22 antibiotics representing five to nine classes (**Supplementary Table 6**).

**Figure 2:**
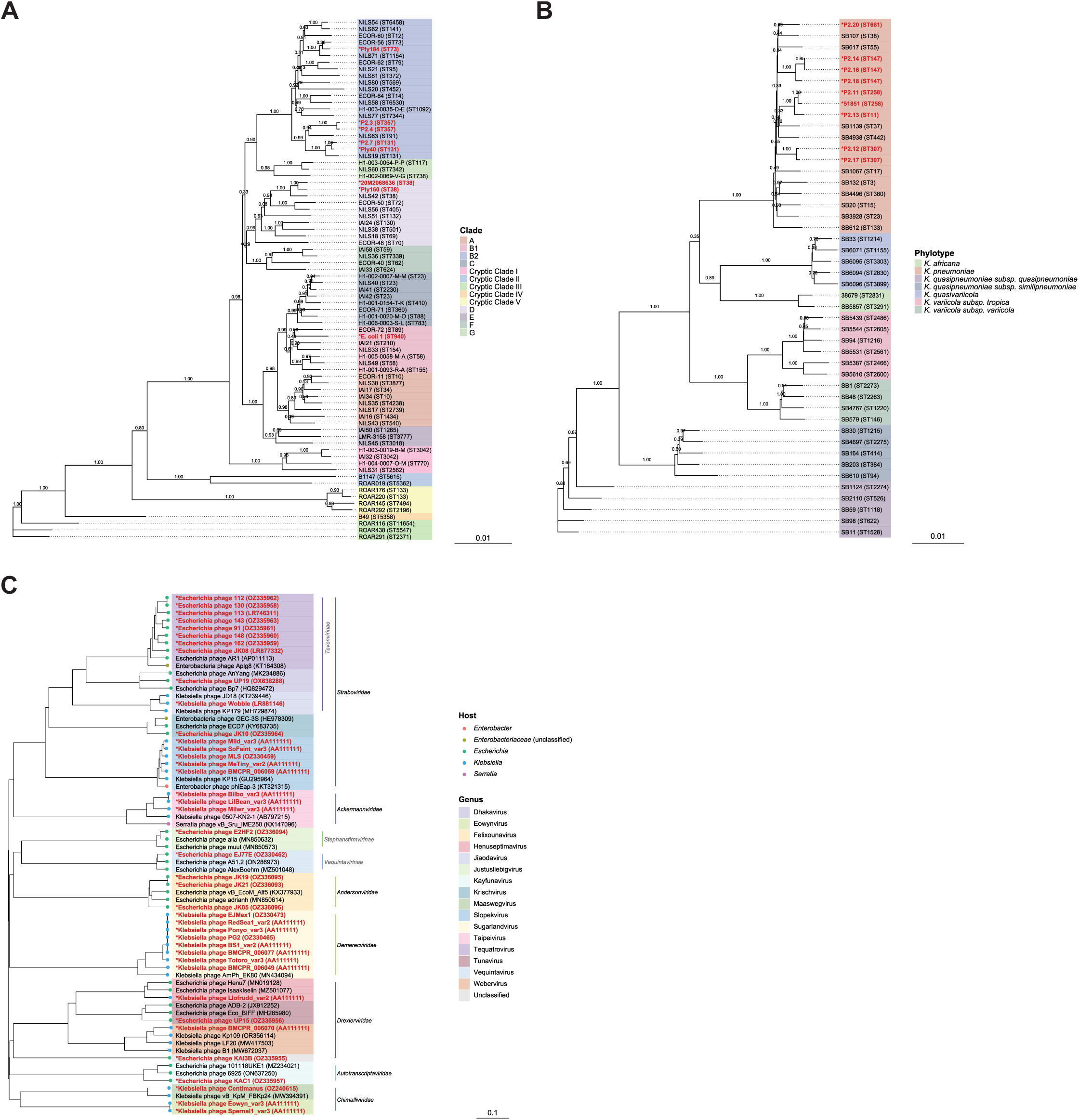
Phylogenetic trees of study phages and isolates with reference strains for context. Trees were generated using distance-based pairwise evolutionary distances^32^ across (A) *E. coli* strains (n = 78) and (B) *Klebsiella* strains (n = 46), and proteome-wide comparisons based on genome-wide sequence similarities computed by tBLASTx^33^ across (C) the *Klebsiella* spp. and *E. coli* phage collection (n = 67). Scale bars refer to the evolutionary distance (number of substitution events per character). Branch confidence values based on the rate of elementary quartets (REQ) are displayed as decimals in the centre of branches. For phylogenetic context, bacterial strains and phages from the current study, indicated in red, were compared to representative collections from the Picard collection^34,35^, Institute Pasteur^36^, and the Virus Metadata Resource (VMR; https://ictv.global/vmr) for *E. coli*, *K. pneumoniae* species complex, and phages, respectively.

Similarly, phages were excluded if they were temperate (n = 1), failed to infect any strain (13; ∼22%), showed low efficacy based on the criteria described in the methods (11; 19%), or could not be amplified to >10 PFU·ml^-1^ (7; ∼12% *Klebsiella* phages; 1 *E. coli* phage).

From the remaining 46 candidates, 43 were selected for downstream analysis: 20 *E. coli* phages and 23 *Klebsiella* spp. phages (**Supplementary Table 7**). The *E. coli* phages represented members of four families (*Andersonviridae*, *Autotranscriptaviridae*, *Drexlerviridae*, and *Straboviridae*) and nine genera (*Dhakavirus*, *Felixounavirus*, *Justusliebigvirus*, *Kayfunavirus*, *Krischvirus*, *Tequatrovirus*, *Tunavirus*, *Vequintavirus*, and a new viral genus). The *Klebsiella* phages were similarly diverse, representing five families (*Ackermannviridae*, *Chimallaviridae*, *Demerecviridae*, *Drexlerviridae*, and *Straboviridae*) and eight genera (*Eowynvirus*, *Henuseptimavirus*, *Jiaodavirus*, *Maaswegvirus*, *Slopekvirus*, *Sugarlandvirus*, *Taipeivirus*, and *Webervirus*) (**Figure 2C**). Three phages (*Escherichia* phages KAH1 and KAL1, *Klebsiella* phage KpSoF) were unable to be sequenced using current technologies (**Supplementary Table 7**). All sequenced genomes lacked AMR determinants and prophage-associated genes, supporting their suitability for therapeutic development.

### Sensititre Plate Analysis and Genomic Insights into Phage-Driven Variability in Antibiotic Interactions and Host Range

To quantify phage-antibiotic effects on bacterial growth, 93 *E. coli*-phage and 52 *K. pneumoniae*-phage combinations (**Supplementary Tables 8** and **9**) were tested using Sensititre GN7F plates, a quality-controlled, commercially available product, comprising 24 different antibiotics, using between 1-7 concentrations per antibiotic (**Supplementary Tables 10** and **11**). For each strain, only phages that resulted in some degree of clearance were included. In total, 13,485 phage-antibiotic interaction data points (**Supplementary Table 12**) were generated, of which ∼17% (n = 2,316) showed significant alterations in bacterial growth (**Supplementary Table 13**).

Phage-antibiotic interactions involving *K. pneumoniae* showed a greater overall impact compared to those with *E. coli* with ∼25% (n = 1,218) and ∼13% of interactions (n = 1,098) respectively, resulting in significant changes to bacterial growth, (χ^2^(1, n = 13,485) = 340.208, *p* < 0.00001). Across the dataset, ∼16% (n = 2,167) of interactions were additive, while ∼1% (n = 149) were antagonistic. Antagonism was rare in both species (*E. coli*, 0.8%, n = 110; *K. pneumoniae*, 0.3%, n = 39), while additive effects were more frequent in *K. pneumoniae* (24%, n = 1,179) than in *E. coli* (11%, n = 988).

Interaction patterns were explored through hierarchical clustering of aggregated interaction matrices, with experimental values averaged within metadata-defined groups, including ST, antibiotic class, and phage taxonomy; this produced heatmaps highlighting clusters of combinations with similar activity patterns (**Figure 3**). A summary of the findings is presented in **Table 1**.

**Figure 3:**
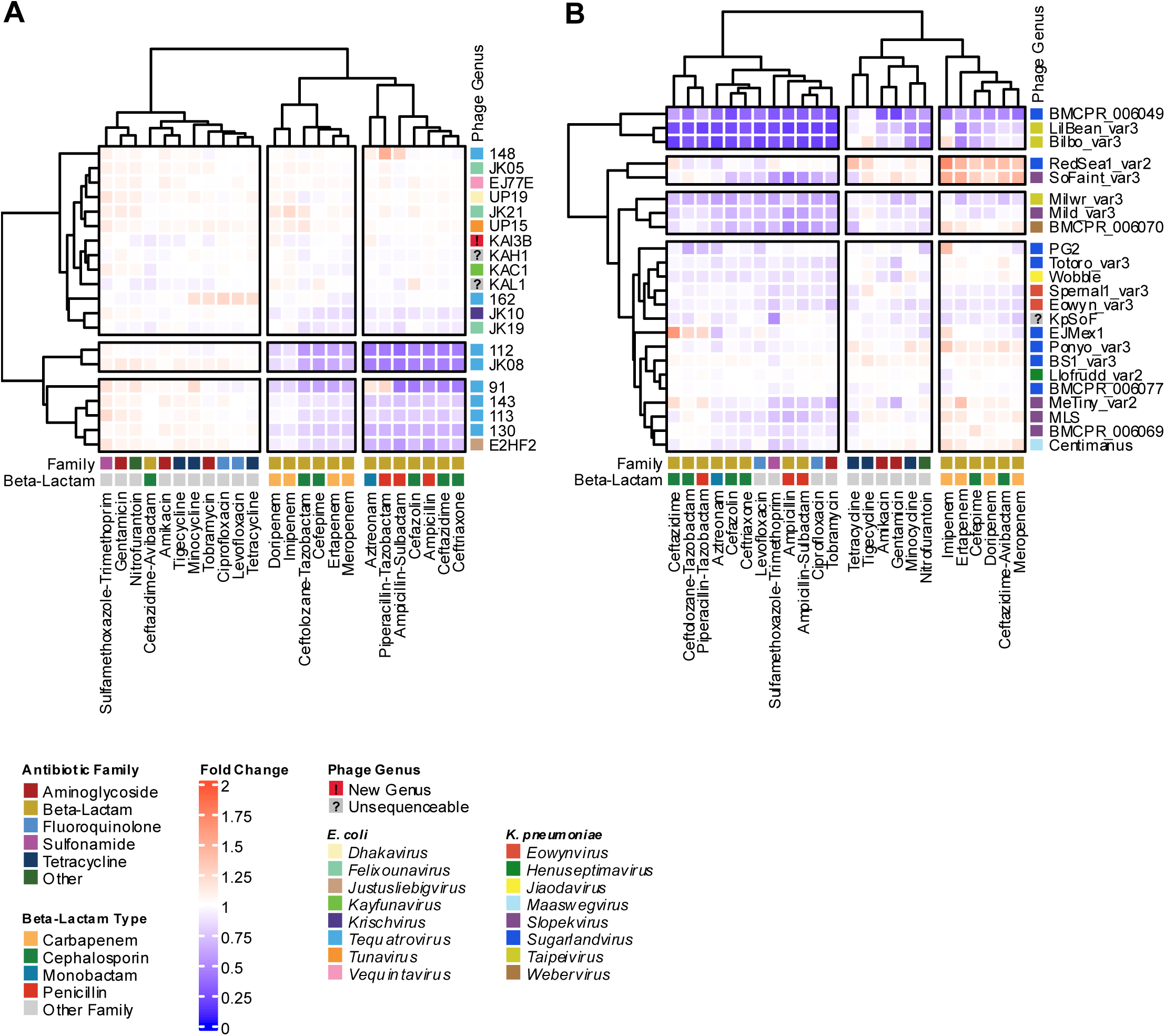
Clustered Heatmap of Phages versus Individual Antibiotics For (A) *E. coli* and (B) *K. pneumoniae*. (A) Clustering the *E. coli* phage-antibiotic interactions based on similarity reveals a distinct 2×2 cluster in the bottom-right corner, representing a group of combined therapies that exhibit an additive effect, specifically β-lactam antibiotics together with *Tequatroviruses*. (B) By contrast, while *K. pneumoniae* interactions reveal a distinct single additive cluster in the top-left corner, there is no consistent association between specific phages or antibiotics, and the additivity appears to be phage-dependent. The rows in the heatmap represent the different phages tested against the isolates, with the coloured square to the left of the phage names indicating the corresponding phage genus. The columns in the heatmap represent the various antibiotic types, with the coloured squares above the antibiotic names indicating the corresponding family (top row), and subtype within the β-lactam family (bottom row). The colour of the squares indicate fold change of bacterial growth following combined phage and antibiotic treatment relative to treatment with the antibiotic alone.

**Table 1:** Key Phage-Antibiotic Interaction Patterns.

| Interaction Context | Phage & Antibiotic | Genomic Similarity | Observation |
| --- | --- | --- | --- |
| **Class Synergy** | All Tequatroviruses + β-lactams (*E. coli*) | >89% ANI | **Strongly Additive.** Consistent enhancement of bacterial killing across the genus. 2 outliers (phages 148 and 162). |
| **Genomic Variants** | *Tequatrovirus* phages 112 vs. 130 (*E. coli*) | >99.9% ANI, differ in 2 genes (each present in only one of the phages) | **Divergent Phenotypes.** The phages displayed an altered host range. 112 had greater additivity with β-lactams, and was additive (vs 130's antagonism) with nitrofurantoin, minocycline, and tobramycin. |
| **Genomic Variants** | Phages JK19 vs. JK21 (*E. coli*) | >99.9% ANI, differ in 5 genes (2 present in each of the phages, and one frameshift/early truncation) | **Opposing Outcomes.**<br>The phages displayed a distinctly altered host range. JK19 was weakly additive while JK21 was weakly antagonistic with β-lactams. JK19 resulted in five decreases and two increases in MIC (one resulting in change to susceptibility), whereas JK21 resulted in two increases in MIC (one resulting in change to susceptibility). |
| **Genomic Variants** | Phages Eowyn_var3 vs. Sernal1_var3 (*K. pneumoniae*) | >99.9% ANI, differ by two insertions in non-coding sequence and two genes, including a tail fibre unique to Eowyn_var3. | **Divergent Phenotypes.** The phages displayed an altered host range. Small differences in additivity intensity. Sernal1_var3 antagonised with gentamicin and tigecycline whereas Eowyn_var3 did not. |
| **Genomic Variants** | Phages LilBean_var3 vs. Bilbo_var3 (*K. pneumoniae*) | >99.9% ANI, differ in four genes (of which two encode tail proteins). | **Divergent Phenotypes.** The phages displayed a slightly altered host range. Bilbo_var3 was antagonistic with imipenem and tigecycline; LilBean_var3 had no effect. |
| **Genus Divergence** | *Sugarlandvirus* genus (*K. pneumoniae*) | >99.5% ANI BMCPR_006077, Ponyo_var3, BS1_var2, PG2, RedSea1_var2, EJMex1<br>Totoro_var3 ~95%<br>BMCPR_006049 ~83% | **Lineage-Specific Effects.** All had distinct host range differences and distinct interaction patterns. Phage BMCPR_006049 (distinct species) was strongly additive with all antibiotics; RedSea1_var2 and Ponyo_var3 antagonised with carbapenems and tetracyclines. EJMex1 antagonised with several β-lactams. BMCPR_006077 and BS1_var2 had no notable interactions at all. |
| **Functional Convergence** | E2HF2 & Tequatroviruses (*E. coli*) | Unrelated | **Similar Interactions.** Both E2HF2 and 112, JK08, 91, 143, 113, 130 all were additive with β-lactams and displayed mild antagonism with other antibiotics, despite being genetically unrelated. |
| **Genus Divergence** | Tequatroviruses (*E. coli*) | >89% ANI | **Divergent Phenotypes.** Phages 112, 113, 130, 143, and JK08 showed remarkably similar interaction patterns. Phage 91 antagonised with aztreonam and piperacillin-tazobactam, unlike the others. Phages 148 and 162 displayed completely different interaction patterns. |
| **Functional Convergence** | RedSea1_var2 & SoFaint_var3 (*K. pneumoniae*) | Unrelated | **Similar Antagonism.** Both were strikingly antagonistic with carbapenems, cefepime, and ceftazidime-avibactam, despite being genetically unrelated. |
| **Unexpected Synergy** | Several *K. pneumoniae* and *E. coli* phages (MeTiny_var2, BMCPR_006049, LilBean_var3, Milwr_var3, BMCPR_006070, Eowyn_var3, and, to a lesser degree, EJMex1, KpSoF, KAL1, and KAI3B) and aminoglycosides and tetracycline | N/A | **Unexpected Additivity.** Phage additivity persisted despite antibiotic inhibition of protein synthesis. |

### E. coli

In *E. coli*, combining the 13 β-lactams with six Tequatroviruses (91, 112, 113, 130, 143 and JK08) produced a strong additive signal (**Figure 3A**). In contrast, other members of the genus diverged from this pattern: phage 148 was antagonistic, whereas phage 162 showed no detectable effect.

All Tequatroviruses shared >89% average nucleotide identity (ANI). Notably, phages 148 and 162, which lacked additive effects, clustered centrally within the genus phylogeny, whereas JK08, the most genetically divergent member, retained strong additive interactions similar to the others (**Supplementary Table 14**). Ceftazidime-avibactam showed no interaction, irrespective of phage background.

Two coliphage pairs (112/130 and JK19/JK21) shared >99.9% ANI yet differed in host range and antibiotic interaction profiles, with the most pronounced differences observed within the latter pair.

Phages 112 and 130 displayed largely similar host ranges; however, five strains were uniquely sensitive to one phage, and they demonstrated differential lytic efficiency. Phage 112 also exhibited greater additivity, with >20% fold change in 9 of the 13 β-lactam antibiotics. Subtle but consistent differences were observed for nitrofurantoin, minocycline and tobramycin, which were additive with phage 112 but antagonistic with phage 130. These phenotypic differences may be associated with two hypothetical genes unique to each genome (**Supplementary Table 14**).

Similarly, Felixounaviruses JK19 and JK21 differed substantially in host range, each infecting eight unique strains, with variable lytic efficiency on two others.

Phage JK19 showed weak but broadly additive interactions with β-lactam antibiotics, while JK21 was weakly antagonistic. Modest fold change differences were observed in 11 of 24 antibiotics, eight of which displayed opposing interaction profiles (cefepime, ceftazidime, ceftolozane-tazobactam, ertapenem, gentamicin, meropenem, nitrofurantoin, and piperacillin-tazobactam).

JK19 increased MICs in two strain-antibiotic combinations, (one altering susceptibility), and decreased MICs in five, whereas JK21 increased MICs in two combinations, (one altering susceptibility). These differences may be associated with a truncated hypothetical gene in JK21 containing two amino acid substitutions, together with four additional hypothetical genes, that differ between the genomes (**Supplementary Table 15**).

Overall, antagonism in *E. coli* combinations was infrequent and modest in magnitude (**Figures 3**, **4** and **5**). Examples include phage 148 with piperacillin-tazobactam, phage 91 with aztreonam and piperacillin-tazobactam, and phage 162 with fluoroquinolones, tetracyclines (minocycline, tetracycline), and tobramycin. This indicates that some phages display cross-class antagonistic tendencies. In contrast, amikacin and tigecycline and phages EJ77E, KAH1, KAI3B, and JK05 produced <10% change in all combinations tested.

**Figure 4:**
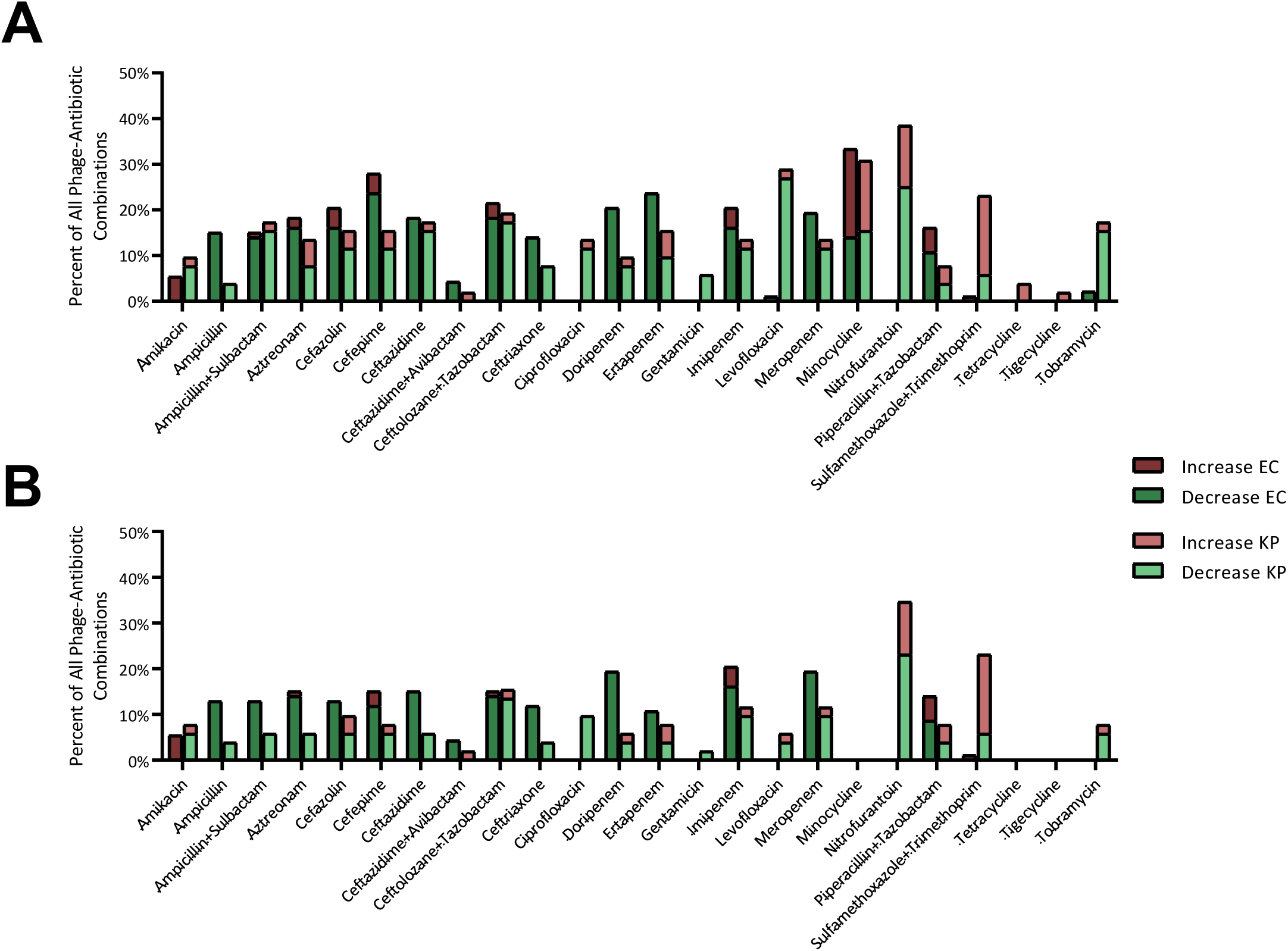
Effects of Dual-Therapy Combinations on Antibiotic MICs and Susceptibility, Per Antibiotic. The percentage of phage-antibiotic combinations, classified by antibiotic type, resulting in changes to (A) MIC and (B) susceptibility. Changes for *E. coli* are represented by the dark bars, and changes for *K. pneumoniae* by the light bars. Red segments indicate the percentage of combinations for each antibiotic resulting in an increased MIC or reduced susceptibility, whereas green segments indicate the percentage of combinations for each antibiotic resulting in a decreased MIC or increased susceptibility.

**Figure 5:**
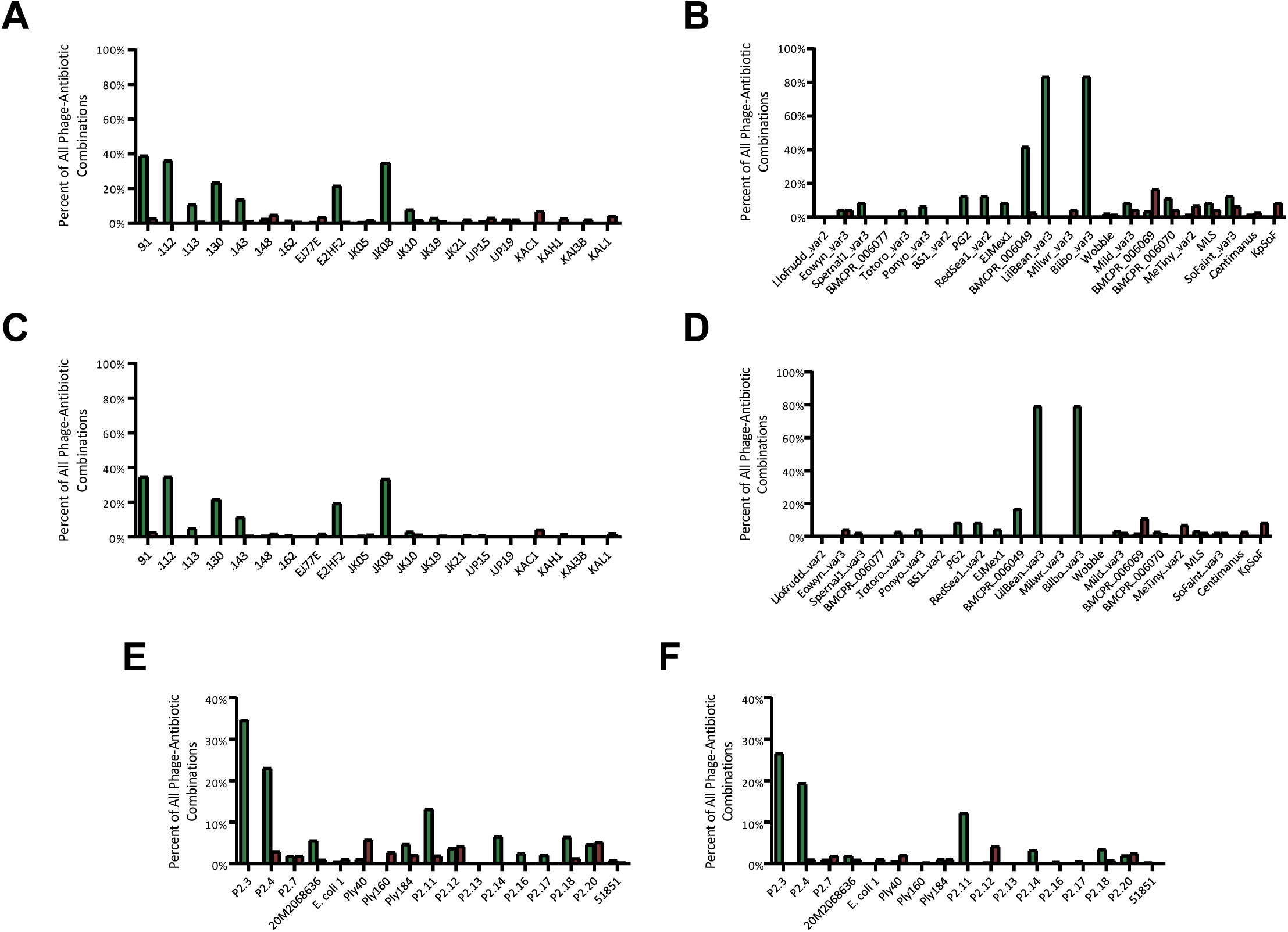
Effects of Dual-Therapy Combinations on Antibiotic MICs and Susceptibility, Per Strain and Phage. The percentage of phage-antibiotic combinations resulting in changes to (**A**) *E. coli* MIC per phage, (**B**) *E. coli* susceptibility per phage, (**C**) *K. pneumoniae* MIC per phage, (**D**) *K. pneumoniae* susceptibility per phage, (**E**) MIC by bacterial strain for both *E. coli* and *K. pneumoniae*, and (**F**) susceptibility by bacterial strain for both species. While not constituting a pattern, significant effects were evident for specific phages and bacterial strains. For example, *Klebsiella* spp. phages LilBean_var3 and Bilbo_var3 increased bacterial susceptibility to ∼80% of antibiotics tested, while *E. coli* phages 91, 112, and JK08 increased susceptibility to ∼35% of antibiotics tested. At the bacterial strain level, *E. coli* strain 2.3 showed increased susceptibility in ∼26% of phage-antibiotic combinations tested, and *E. coli* strain 2.4 and *K. pneumoniae* strain 2.11 showed increased susceptibility in ∼20% and ∼15% of combinations, respectively.

### K. pneumoniae

Phage-antibiotic interaction profiling revealed substantial variation in additive and antagonistic effects across *Klebsiella* phage genera compared with those observed in *E. coli* (**Figure 3B**). Strong additive effects with nearly all antibiotics tested, particularly β-lactams and fluoroquinolones, were observed in six phages: BMCPR_006049 (*Sugarlandvirus*), LilBean_var3 and Bilbo_var3 (*Taipeivirus*), and, to a lesser extent, Milwr_var3 (*Taipeivirus*), Mild_var2 (*Slopekvirus*), and BMCPR_006070 (*Webervirus*). Antagonistic interactions were generally low-intensity, and restricted to carbapenems and ceftazidime-avibactam when combined with *Sugarlandvirus* phages Ponyo_var3, RedSea1_var2, and EJMex1, as well as several *Slopekvirus* and *Maaswegvirus* phages.

Notably, pronounced phenotypic divergence was observed among the members of the genus *Sugarlandvirus* (**Supplementary Table 16**). BMCPR_006049 exhibited one of the highest degrees of additivity, enhancing the activity of 96% (n = 23) of antibiotics across all classes with the exception of tigecycline, for which no significant change in bacterial growth was observed. In contrast, RedSea1_var2 and Ponyo_var3 displayed pronounced antagonism across multiple antibiotic classes, particularly β-lactams and tetracyclines. Genomic analysis revealed that both BMCPR_006049 and Totoro_var3 represent distinct species-level lineages within the genus *Sugarlandvirus* (*Sugarlandvirus storm* at ∼83% ANI, and *Sugarlandvirus totoro* at ∼95% ANI), while the remaining six, including RedSea1_var2 and Ponyo_var3, belong to a highly similar cluster sharing >99.5% ANI, highlighting that intragenus variation in phage-antibiotic interactions is substantial.

Intraspecies variation was also evident, though less pronounced, within the group of six highly similar phages. Phage PG2 exhibited additive interactions to different antibiotics more frequently than the others. In contrast, antagonism with carbapenems was more pronounced for RedSea1_var2 and Ponyo_var3, and antagonism with ceftazidime was observed only in EJMex1. These differences in antibiotic interaction profiles may be linked to the minor genomic variations observed between the phages.

It should be noted that EJMex1 and PG2 are identical (0 single nucleotide polymorphisms, or SNPs); nonetheless, their interactions with antibiotics differ. The strain used in these tests, *K. pneumoniae* P2.18, displays a degree of heteroresistance (where subpopulations within a culture display differing levels of antimicrobial resistance^37^). The existence of such subpopulations may contribute to the observed phenotypic differences despite the underlying genetic identity.

Variability at the species level was observed within the genus *Eowynvirus*. While both Spernal1_var3 and Eowyn_var3 displayed mild additive effects across the range of antibiotics tested, Spernal1_var3 differed in that it displayed weak antagonism with tigecycline and gentamicin. Despite the high overall similarity (>99.99% ANI), the two phages display limited, localised genomic differences. These include a substitution in a hypothetical protein as well as a tail fibre-associated gene unique to Eowyn_var3 (**Supplemental Table 17**).

More pronounced species-level variation was observed within the genus *Taipeivirus* (**Supplementary Table 18**). While LilBean_var3 had no impact on the efficacy of imipenem and tigecycline, Bilbo_var3 showed some degree of antagonism. Despite sharing >99.99% ANI, these phages differ by several single-residue substitutions and deletions, including in genes encoding a major head protein, a tail protein, and a tail fibre protein, overall suggesting that even minor genomic changes can drive distinct phenotypic outcomes.

### Impact of Phage-Antibiotic Combination on MIC and Clinical Susceptibility

Sensititre plate assays were used to assess the impact of phage-antibiotic combinations on each antibiotic’s minimum inhibitory concentration (MIC).

Antibiograms for each isolate (**Supplementary Table 6**) were compared with corresponding phage-antibiotic combinations (**Supplementary Table 19**). Of the 3,480 phage-antibiotic combinations, ∼13% (456 combinations) resulted in a change in MIC, and ∼8.4% (292) changed susceptibility classification.

For MIC shifts, bacterial species were significantly associated with outcome (increase, decrease or no change) (χ²(2, n = 3,480) = 10.72, *p* = 0.005). Among combinations with altered MICs, the direction of change also differed between species (χ²(1, n = 456) = 7.81, *p* = 0.005).

Similarly, susceptibility changes were significantly associated with species overall (χ²(1, n = 3,480) = 17.16, *p* = 0.0002), and the direction of change among affected combinations differed significantly (χ²(2, n = 456) = 17.22, *p* = 0.00003).

A general trend toward reduced antibiotic MICs was observed for *E. coli* phage-antibiotic interactions, with 10% (230) of combinations resulting in decreased MICs and 2% (47) resulting in increased MICs (**Figure 4A**). When translated into clinical susceptibility, ∼8% (171) of combinations resulted in increased susceptibility and >1% (20) resulted in decreased susceptibility. A similar observation was found for *K. pneumoniae,* with ∼10% (129) of combinations resulting in decreased MIC and ∼4% (50) resulting in increased MICs. When translated into clinical susceptibility, ∼6% (71) of combinations resulted in increased susceptibility and ∼2% (30) resulted in decreased susceptibility (**Figure 4B**).

For both *E. coli* and *K. pneumoniae,* phage addition consistently reduced MICs for ampicillin and ceftriaxone, and reductions predominated across all but one β-lactam antibiotic.

In contrast, increased MICs were observed for the tetracyclines tetracycline and and tigecycline. Species-specific differences were also evident; in *E. coli*, MICs remained unchanged for ciprofloxacin, gentamicin, nitrofurantoin, tetracycline, or tigecycline, whereas increases were observed for amikacin and sulfamethoxazole-trimethoprim. In *E. coli*, adding phage to ceftazidime-avibactam decreased the MIC, in contrast to the increase found with *K. pneumoniae*.

While patterns in MIC could be observed between and across *E. coli* and *K. pneumoniae,* there were also observed changes in MIC which were both phage- and bacterial strain-dependent (**Figure 5**). Of the 20 coliphages, five phages (25%; JK21, KAC1, KAH1, KAI3B, and KAL1) were only associated with an increased MIC, with three phages (15%; 112, 162, JK08) associated with a decreased MIC (**Figure 5A-B**). In contrast, with *K. pneumoniae* phages, only two of the 23 phages (∼9%; Milwr_var3 and KpSoF) were associated with an increased MIC (**Figure 5C-D**). A higher proportion of phages were associated with a decreased MIC, with nine phages (∼39%; Eowyn_var3, Spernal1_var3, Totoro_var3, Ponyo_var3, PG2, RedSea1_var2, EJMex1, LilBean_var3, and Bilbo_var3) associated with this decrease. There were also three *K. pneumoniae* phages (∼13%; Llofrudd_var2, BMCPR_006077, and BS1_var2) that never induced changes to MIC. Within the *E. coli* strains tested, strain Ply160 only exhibited increased MIC in combination with a phage, if a change was observed (**Figure 5E**). In contrast strain 2.3 only exhibited a decreased MIC. *K. pneumoniae* strains 2.16 and 2.17 exhibited MIC decreases exclusively (**Figure 5F**).

Finally, while *K. pneumoniae* exhibits interactions with all the antibiotics tested, it also accounts for the majority of the antagonistic effects; in contrast, *E. coli* displays predominantly additive interactions, with the exception of minocycline, but also has no interactions with four antibiotics (ciprofloxacin, gentamicin, nitrofurantoin, and tigecycline). This comparison becomes apparent when the interactions are categorised by changes in MIC and susceptibility, highlighting species-specific patterns (**Figure 6**).

**Figure 6:**
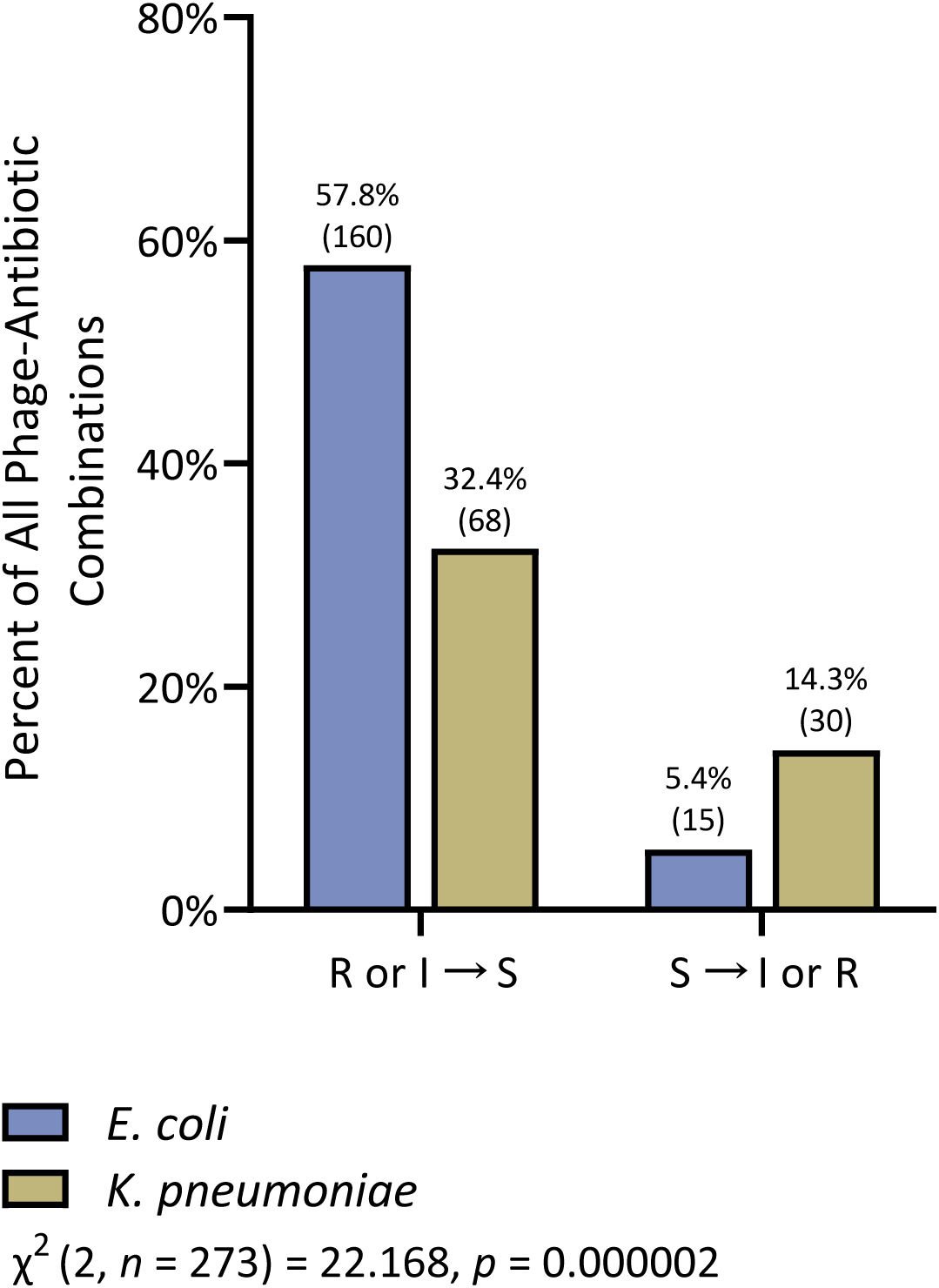
Percent of Phage-Antibiotic Combinations Resulting in Gain and Loss of Susceptibility. Phage-antibiotic combinations were classified according to their observed effects on bacterial MIC and antibiotic susceptibility. Combinations were grouped into categories reflecting either development of susceptibility or, conversely, development of resistance, according to clinical breakpoints. Interpretation followed EUCAST guidelines v16.0^38^, whereby R indicates Resistant, S indicates Susceptible, and I indicates Susceptible, Increased Exposure. The percentage of combinations within each category is presented, with the total number indicated above each column to facilitate comparison between *E. coli* and *K. pneumoniae*. Across both species, development of susceptibility accounted for the most frequent interaction type. However, the next most frequent type of interactions differed between species: in *E. coli*, this involved reductions in MIC without a corresponding change in susceptibility, whereas in *K. pneumoniae* this involved the development of resistance. A chi-square test of independence demonstrated a statistically significant association between interaction direction and bacterial species (*p* < 0.00001), indicating that the distribution of interaction outcomes differed between *E. coli* and *K. pneumoniae*.

## Discussion

Previous phage-antibiotic studies have used small panels of strains and phages and antibiotics^7–17^, which limits their generalisability. To address this, our high-throughput approach allowed us to assess a broad range of interactions, which enabled a statistical analysis of trends and potential links between phage genetics and interaction type. These data provide a rational basis for prioritising phage-antibiotic combinations for subsequent mechanistic investigation.

Commercially available Sensititre GN7F plates, designed for MIC determination in Gram-negative pathogens, were used to assess *E. coli* and *K. pneumoniae*, two leading causes of urinary tract infections and the primary focus of this study. The plates comprise 24 clinically relevant antibiotics, each tested across multiple concentrations, enabling the simultaneous assessment of 93 antibiotic conditions per plate. This configuration provided a robust high-throughput platform to detect MIC shifts and changes in susceptibility following phage addition. Importantly, the use of a validated and standardised system ensured rapid implementation and high reproducibility, and minimised the variability associated with bespoke assay formats. As a widely used, regulatory-approved diagnostic platform, Sensititre provides a practical foundation for future clinical phage-antibiotic susceptibility testing.

A limitation of this study is that the antibiotic concentrations tested do not reflect *in vivo* pharmacokinetics or urinary or plasma exposures; for example, peak urinary concentrations of ciprofloxacin (∼200 µg/ml^39,40^) substantially exceed those represented on the plate. Furthermore, clinical infections involve complex host-pathogen-drug interactions that are not captured *in vitro*. However, the observed MIC shifts demonstrate that phages can enhance antibiotic activity and alter susceptibility profiles, which suggests that they have the potential to restore efficacy of compromised drugs.

Across conditions, additive phage-antibiotic interactions were markedly more frequent than antagonistic ones, with no instances of pronounced antagonism, defined here as a >2-fold increase in bacterial growth following phage addition. This is of particular interest given that a quarter of the antibiotics tested, e.g. aminoglycosides and tetracyclines, work by inhibiting protein synthesis, and nitrofurantoin induces multifaceted damage to cellular processes essential for phage replication **(Supplementary Table 11**). Despite these potentially inhibitory mechanisms, phage activity was largely preserved and, in some cases, enhanced in combination treatments.

Overall, 17.0% of the phage-antibiotic interactions were statistically significant, with clear species-specific patterns. Significant interactions were nearly twice as frequent in *K. pneumoniae* than in *E. coli* (25.2% vs. 12.7%) and were more often additive (96.8% vs. 90.0%). Chi-squared analyses confirmed significant associations between bacterial species and the frequency and direction of MIC and susceptibility changes, which highlights the importance of incorporating species-specific effects into future pharmacokinetics/pharmacodynamics-informed and *in vivo* studies.

The trends observed likely reflect differences in intrinsic antibiotic resistance and phage susceptibility between strains. The *K. pneumoniae* panel had a higher baseline resistance, which provided a broader dynamic range where phage-mediated effects could be seen. Together, our findings indicate that both the magnitude and direction of phage-antibiotic interactions depend on the antibiotic class and mechanism of action and on the underlying resistance landscape of the bacterial host.

Within the *E. coli* dataset, interactions with β-lactam antibiotics were predominantly additive, consistent with previous reports in both *E. coli* and *K. pneumoniae*^20,23,41–48^. This supports broader observations that β-lactams often enhance phage activity, potentially by altering cell wall architecture in ways that facilitate phage adsorption or release. Taken together, these data support a model in which certain phages act less as standalone antimicrobials and more as functional enhancers of β-lactam activity. Rather than replacing antibiotics, these phages appear to shift bacterial susceptibility in ways that restore or extend the efficacy of existing β-lactams, including in resistant strain backgrounds.

Notably, several Tequatroviruses displayed highly similar additive interaction profiles across antibiotics. While some of these were closely related or taxonomically identical, others with comparable genetic distances showed distinct host ranges and more antagonistic patterns, indicating that genetic relatedness alone does not explain interaction behaviour. A similar additive profile was also observed for phage E2HF2, a *Justusliebigvirus*, suggesting that such responses are not confined to a single lineage. Given the genetic diversity of Tequatroviruses, the consistency observed here likely reflects functional convergence rather than genus-wide behaviour, highlighting the need for further work to define the genetic and mechanistic drivers of these interactions.

Importantly, this β-lactam-supporting phenotype was not evenly distributed across the phage collection but was strongly enriched among a subset of Tequatroviruses, which displayed consistent additive interactions across multiple β-lactam subclasses.

This recurrent pattern suggests a lineage-linked functional capacity for β-lactam potentiation, while the presence of both additive and non-additive phenotypes within the genus highlights that this is not a universal property of the genus Tequatrovirus.

In *K. pneumoniae*, clustering analysis revealed partially distinct phage groupings, although these were less clearly resolved than those observed for *E. coli*. The two phages LilBean_var3 and Bilbo_var3, which form a new species in the genus *Taipeivirus*, showed broadly additive interactions across a wide range of antibiotics. A third *Taipeivirus*, Milwr_var3, exhibited a similar but weaker pattern, consistent with conserved functional traits within this genus.

Several phages (MeTiny_var2, BMCPR_006049, LilBean_var3, Milwr_var3, BMCPR_006070, Eowyn_var3, and, to a lesser degree, EJMex1, KpSoF, KAL1, and KAI3B) had a distinctive interaction profile, characterised by consistently additive effects with aminoglycosides and tetracyclines across all tested doses. Although pronounced additive or antagonistic effects were largely absent, this selective compatibility with protein synthesis inhibitors (that are typically expected to limit phage replication) is notable and suggests a phage-specific resilience to host translational inhibition.

Eight Sugarlandviruses were analysed, which consist of six highly similar phages (BMCPR_006077, Ponyo_var3, BS1_var2, PG2, RedSea1_var2, and EJMex1; >99.82% ANI) and two phages from other species (*Sugarlandvirus totoro* - Totoro_var3, and *Sugarlandvirus storm* - BMCPR_006049). Despite their high genomic similarity, the six phages showed reproducible differences in terms of their host range and antibiotic interaction profiles. These differences are likely driven by variation across 14 genes, including two encoding distinct tail proteins. Although additional polymorphisms were present, structural variation in adsorption-associated proteins is the most plausible driver of the observed phenotypic divergence.

An interesting case of functional convergence was seen between RedSea1_var2 (*Sugarlandvirus*) and SoFaint_var3 (*Slopekvirus*), Despite being taxonomically unrelated and sharing no detectable genomic similarity, these phages displayed strikingly similar interaction profiles, characterised by consistently low antagonism in the presence of carbapenem antibiotics.The mechanistic basis for this convergence remains unclear. Known phage-carbapenem interactions, such as those involving modulation of the MexXY-OprM efflux system that mediates meropenem entry in *Pseudomonas aeruginosa*^50–52^ have not been described in *Klebsiella* spp., suggesting that alternative or indirect mechanisms may be involved.

Several phages (KAI3B, JK05, EJ77E, KAH1, Llofrudd_var2, and BMCPR_006077) showed no measurable phage-antibiotic interactions across the conditions tested. While these combinations did not confer synergistic or additive benefits, the absence of antagonism is itself notable.These phages did not impair antibiotic activity, indicating that they could be incorporated into broader or empirically designed phage cocktails without compromising concurrent antimicrobial therapy.

Reported phage-antibiotic interactions in the literature vary widely, reflecting substantial differences in experimental systems, endpoints, and analytical frameworks. Approaches have included plaque assays, CFU enumeration, optical density-based growth curves, animal survival models, biofilm assays, synograms, and shifts in MIC or MBEC values^20,23,41,44,45,48,53–58^.

Even for the same antibiotic or phage genus, outcomes are frequently inconsistent. For example, studies combining *E. coli* phages with ciprofloxacin^44,45,52,59^ or kanamycin^27,41,43–45^, and studies combining *Klebsiella* phages with ceftazidime^23,48,49,60^ have variably reported synergy, additivity, antagonism, or no effect. These discrepancies are likely to be driven by differences in bacterial strain-phage pairing, antibiotic dosing, and experimental design, that limits direct comparability between previous studies and with the work described here.

Despite this variability, our results match broader patterns which have been reported in multiple studies. Tequatroviruses have generally shown positive or neutral interactions with ampicillin^41,55,61^, meropenem^62^, and ceftazidime^45^. Mixed outcomes have been reported for tetracycline and ciprofloxacin, where both negative and positive, and positive and neutral effects occur, respectively^41,45,55,61,63,64^. Similarly, Vequintaviruses show neutral interactions with ampicillin^55^, Slopekviruses with gentamicin produce synergy^57^ and Jiaodaviruses with gentamicin show antagonism^65^.

Notably, only ten studies to date have examined phage-driven changes in MIC, and just four involved *E. coli* or *K. pneumoniae*^20,66,67^. Although limited, most reported reductions in MIC, with approximately half documenting a shift in susceptibility category.

In contrast to the fragmented and largely small-scale literature to date, therefore, this study represents the first large-scale, systematic analysis of phage-antibiotic interactions spanning multiple bacterial species, phage genera, and antibiotic classes. By interrogating thousands of combinations under controlled and standardised conditions, we demonstrate that phage identity, antibiotic class, and bacterial strain background each critically shape interaction outcomes. In doing so, these data provide a coherent explanation for the inconsistencies reported in earlier studies and begin to define reproducible patterns of additivity, antagonism, and neutrality.

Although the number of strains decreased during the screening process and not all phage–antibiotic combinations were tested across all strains, which may introduce some unintended bias, these limitations were primarily due to practical constraints. Despite this, the study remains to the best of our knowledge the largest of its kind to date, and consistent patterns were still identifiable within the dataset, fulfilling the primary objective of systematically characterising phage–antibiotic interactions.

By moving beyond anecdotal observations, this work establishes a robust and reproducible framework for the rational design of phage-antibiotic combination therapies. Importantly, it demonstrates that an already implemented clinical tool can be leveraged to systematically identify optimised therapeutic pairings. The consistent positive interactions observed between β-lactam antibiotics and Tequatroviruses highlight a particularly promising direction for translational development. Such combinations are particularly attractive from a clinical and regulatory perspective, as they build on well-established β-lactam therapies and suggest a route to revitalise existing antibiotics rather than introducing entirely new antimicrobial classes. Future studies should build on this foundation by integrating genomic, transcriptomic, and phenotypic datasets to resolve the mechanistic basis of these interactions, and by validating key findings in relevant *in vivo* infection models. Together, these approaches will support the development of rational, evidence-based phage-antibiotic therapies to help restore treatment options against multidrug-resistant infections.

## Methods

### Bacterial Isolates and Phages

#### Strains and growth conditions

Bacterial strains used in this study are listed in **Supplementary Table 1**. Bacteria were routinely grown on LB 1.5% (w/v) agar plates (Melford Laboratories Ltd., Ipswich, UK) supplemented with 1 mM CaCl_2_ and incubated overnight (18 h) at 37°C. Liquid cultures were prepared by inoculating a single colony into 5 ml of LB broth and incubating overnight at 37°C with orbital shaking (200 rpm).

#### Phage Isolation and Propagation

Four phages (KAC1, KAI3B, KAL1, and KAL3) were isolated from sewage on *E. coli* using a modified version of a previously described method^68,69^. Briefly, wastewater was collected from the combined sewage influent tank at Minworth Sewage Treatment Works (Birmingham, UK), clarified through a cheesecloth, and then filtered through 0.45 μm pore sized PES filters (Sarstedt, Nümbrecht, Germany). Filtered sewage (5 ml) was enriched with 5 ml LB broth (Melford Laboratories Ltd., Ipswich, UK) supplemented with 1 mM CaCl_2_, combined with 500 μl overnight *E. coli* culture, and incubated overnight at 37°C with orbital shaking. The enriched cultures were centrifuged (8162 × *g*, 10 min) and filtered through 0.2 μm pore sized PES filters (Sarstedt, Nümbrecht, Germany).

Phage presence was assessed by plaque assays: 100 μl of phage (ten-fold dilutions) were combined with 200 μl of overnight target bacterial culture, and 8 ml of LB top agar (0.7% w/v), poured on top of an LB agar plate, and then incubated overnight at 37°C. Following incubation, plates were inspected for the presence of plaques indicative of phage. To obtain clonal phage populations, plaques were purified by repeated isolation streaking on double-layer agar until plaque morphology was uniform. Purified phages were recovered by excising individual plaques, diffusing them in SM buffer for 24 h at room temperature, and filtering through 0.2 μm PES filters (Sarstedt, Nümbrecht, Germany).

Phages and phage host strains used in this study are listed in **Supplementary Table 2** and **Supplementary Table 20**, respectively. To propagate the phages, an overnight culture of the host strain was subcultured 1:60 into 5 ml of LB medium supplemented with 25 mM CaCl_2_ and incubated at 37°C with orbital shaking until the optical density (OD_600_) reached 0.2. At that point, 100 μL of phage stock (1×10^9^ PFU·ml^-1^) was added, and the culture was further incubated overnight at 37°C with orbital shaking (200 rpm). The resulting lysate was centrifuged at 8162 × *g* for 10 min and filtered through 0.2 μm pore size PES filters (Sarstedt, Nümbrecht, Germany), and the PFU·ml^-1^ concentration of the phages was calculated by plaque assays serial dilution.

### Selection of Antibiotic-Resistant and Phage-Sensitive Bacterial Strains

#### Bacterial Screening

Bacterial isolates without documented antibiograms were analysed for the presence of antimicrobial resistance genes using The Comprehensive Antibiotic Resistance Database (CARD)^70^. Isolates that were classified as a MDR or less^71^ or had AMR genes indicating such a classification were excluded from further testing.

#### Phage Host Range Analysis and Efficacy

The spectrum of activity of the phage collection was first assessed using the agar-based spot test assay^72^. The efficacy of infection on solid agar was ranked on a scale of 0-3, corresponding to no observable clearance [0], low clearance [1], moderate clearance [2], and complete clearance [3] (**Supplementary Figure 6**). Combinations scoring 0 were excluded from further analyses, while the remainder of the collection were assessed using liquid medium-based assay designed to emulate the conditions of the Sensititre test, with the aim of identifying the most suitable phages for the experiment.

For the liquid medium-based assay, blank 96 well plates were prepared and inoculated in triplicate using the methods outlined in the manufacturer’s instructions for the GN7F Sensititre plates^73^, and 11 μl of phage stock at a titre of 1×10^8^ PFU·ml^-1^ was added along with the bacteria to the dispensing tube of Mueller-Hinton broth.

Bacterial strains that had low overall phage efficacy (average score <1 for *E. coli* or <0.15 for *Klebsiella* spp. and less than ten effective phages) and phages with low infectivity (average score <0.33) or which failed to reach a titre of at least 10^8^ PFU·ml^-1^ were excluded from further testing.

### Modified Sensititre Plate Protocol

#### GN7F Plate Inoculation

The GN7F plates (**Supplementary Table 11**) (Thermo Scientific, Waltham, MA USA) were prepared and inoculated in triplicate using the methods according to manufacturer’s instructions^73^. For each phage tested in the Sensititre system, 11 μl of phage stock at a titre of 1×10^8^ PFU·ml^-1^ was added along with the bacteria to the dispensing tube of Mueller-Hinton broth, resulting in final concentrations of ∼1×10^5^ for both bacteria and phage (MOI of 1). For control plates, 11 μl of SM Buffer was used.

#### Quantitative Analysis

To measure the changes in bacterial growth under the various phage/antibiotic combination conditions, the size and opacity of the pellet formed at the bottom of each well was measured using ImageJ (Fiji version 1.54f)^74^. Briefly, all images were converted to 32-bit B/W images, the colours inverted, and the background (empty space around the pellets) subtracted (rolling ball radius 50 px, sliding paraboloid, smoothing disabled) to ensure no noise around the pellets.

Regions of interest (ROIs) were positioned around each pellet and the intensity values measured using a custom-written Jython script (**Supplementary Code 1**). Values measured on an empty Sensititre plate with Mueller-Hinton broth served as a blank for normalisation of measured values.

After subtraction of the blank intensity values, the mean intensity values were calculated for each plate triplicate, and the mean intensity values for each well in the phage experiments were normalised against the control values for the relevant strain, such that the values now indicate fold change in relative intensity; i.e., a multiplicative factor where 1 indicates 1× or 100%, values above 1 indicate an increase in bacterial growth, and values below 1 indicate a decrease. Values below 1 were classified as additive, as the experimental design did not permit formal calculation of synergistic effects.

#### Statistical Analysis

Statistical significance was determined using GraphPad Prism 10.4.0.621 by means of 2-way ANOVA with multiple comparisons. For each strain, each phage-antibiotic combination was compared to the control, with the p-value for significance set to 0.05. In order to control the false discovery rate (FDR), or increased probability of false positives associated with multiple comparisons, the Benjamini, Krieger, and Yekutieli two-stage step-up method^75^ was utilised.

#### Clustering and Heatmaps

The experimental data was organised into interaction matrices based on metadata such as antibiotic class, antibiotic mechanism of action, multilocus sequence typing (MLST) of the bacterial strains, and phage taxonomy using Tableau 2024.1.1. For each metadata group for example, all carbapenem-related entries per phage data points were averaged to generate representative values, enabling more comprehensive analyses and revealing patterns not evident in the dataset’s original row-column structure, such as trends across bacterial strains or among individual phages. The results were then arranged using complete linkage clustering based on a Euclidean distance matrix. Heatmaps were generated using R 4.3.1 and RStudio 2023.03.1, using the package ComplexHeatmap 2.15.4.

### Genomic Analysis

#### Genomic DNA Extraction

Genomic bacterial DNA was isolated using the GenElute Bacterial Genomic DNA Kit (Sigma-Aldrich, St. Louis, MO USA), according to the manufacturer’s instructions.

Genomic phage DNA was extracted from liquid, cell-free lysate using a modified phenol:chloroform method^76^. Briefly, phage lysate (1 ml) was digested with 500 μg/mL proteinase K at 65°C for 30 min. The digested lysate was mixed with equal volume of phenol:chloroform:isoamyl alcohol (25:24:1), vortexed, incubated at room temperature for 2 min, vortexed again, and centrifuged (13,000 × *g*, 5 min). The upper aqueous phase was collected, and the process repeated 3-5 times, or until no interface remained between the aqueous and organic phases. The final aqueous phase was transferred to a fresh tube and mixed with one-tenth volume of 3 M sodium acetate (pH 5.2) and 2 volumes of ice-cold 100% ethanol. The mixture was incubated (4°C, 1 h) to precipitate the DNA, then centrifuged (13,000 × *g*, 30 min, 4°C). The supernatant was discarded and the DNA pellet washed with 1 ml of ice-cold 70% ethanol, incubated on ice for 30 min, and centrifuged again. The DNA was then resuspended in 30 μl of ultra-pure, nuclease-free water and quantified using a Qubit fluorometer with the high sensitivity kit (Life Technologies, Carlsbad, CA USA).

Bacterial DNA samples were sequenced by SeqCenter (Pittsburgh, PA USA) using Illumina (2* 150 bp) with libraries prepared with Nextera tagmentation. Phage DNA samples were sequenced in-house using Oxford Nanopore Technology (ONT) Rapid sequencing DNA V14 - barcoding kit (SQK-RBK114.24 or SQK-RBK114.96) on a MinION Mk1c/Mk1B or GridION, with R10.4.1 flow cells^77^.

#### Bacterial Genome Assembly, Annotation, and Phylogeny

Sequencing reads were reduced to a depth of 100 x and assembled *de novo* using Shovill v0.9.0 (https://github.com/tseemann/shovill). Bacterial multi-locus sequence typing (MLST) was carried out via Pathogenwatch v23.3.0, using PubMLST^78^ and EnteroBase^79^. *E. coli* and *Klebsiella* spp. strains were phylotyped using the Clermont method via EzClermont^80,81^ and core genome MLST-based Life Identification Numbers (LIN) codes, respectively.

Phylogenetic analysis was carried out based on distance-based pairwise evolutionary distances using JolyTree 2.1^32^ with pairwise dissimilarity between the genomes estimated using Mash v2.3 ^83^ and branch confidence values generated based on the rate of elementary quartets (REQ)^84^.

The phylogenetic tree was visualised and annotated using GrapeTree^85^ and the R packages R 4.3.1 and RStudio 2023.03.1, using the packages ggtree 3.14.0 and ggtreeExtra 1.19.0^86,87^. For phylogenetic context, representative strains from the Picard^35,88^ and Institute Pasteur collections^36^ were included in the analysis.

#### Phage Genome Assembly, Annotation, Phylogeny, and Analysis

Phage genomes sequenced with Illumina were assembled using a previously described protocol^89^, with minor modifications. Briefly, reads were trimmed with trim_galore, and assembled *de novo* with SPAdes v3.15.5 “only-assembler”^90^, and genomes were then trimmed to remove repeats introduced for circular genomes by SPAdes.

Prior to use, phages that were acquired from collaborators were resequenced using the *Plaque-2-seq* approach^77^. Briefly, phages were spotted onto their respective host bacterial lawns and incubated overnight to confirm their activity. A soft agar plug was collected from the middle of the resultant clearing zone and transferred to a 1.5 mL Eppendorf tube containing 100 μL SM buffer, vortexed and left for 1 hour for phages to disperse from the agar. These samples underwent DNase I (Thermo Scientific) treatment, followed by incubation at 95°C for 10 minutes.

Amplification was achieved using EquiPhi29™ DNA Amplification Kit (Thermo Scientific) following the manufacturer’s guidelines, with DNA debranched using S1 nuclease (Thermo Scientific). Amplified DNA was sequenced using ONT. Sequencing libraries were prepared using Rapid sequencing DNA V14 - barcoding kit (SQK-RBK114.96), and sequenced on a GridION, with R10.4.1 flow cells. Basecalling using the High-accuracy v4.3.0 - 400 bps model and subsequent trimming of the reads of barcodes and adapter sequences were performed with MinKnow.

Long reads were converted to short reads of length 300 bp with a python script, followed by normalisation with bbnorm “--target=150” and the assembly with SPAdes v3.15.5 “-s --only-assembler”^91^.

All bacteriophage genomes were annotated with Prokka v1.14.6^92^ using PHROGS HMMs^93^ and reorientated with the end of the first intergenic region upstream of the terminase large subunit (TerL) being used as the “start” of the phage genomic sequence. All genomes were submitted to the European Nucleotide Archive (ENA) (see supplementary table for the accession numbers). Phages were classified into existing genera and species using *taxMyPhage* v0.3.6 with ”run” setting^94^.

DNAdiff (part of MUMmer 4.0.0 rc1)^95^ was employed to identify single nucleotide polymorphisms (SNPs) between phages, and Roary 3.13.0^96^ to define core genome differences, which were compared using EMBOSS Needle (EBLOSUM62 matrix, 10.0 gap penalty, 0.5 extend penalty)^97^. Phylogenetic analysis of phage genomes generated in this study was carried out with ViPTreeGen 1.1.3^33^, with the addition of representative species of defined phage families^98^.

## Funding Acknowledgement

This work was supported by the ALICE High Performance Computing facility at the University of Leicester, and UKRI grants APP63478 (JPIAMR) and BB/Y51374X/1 (BBSRC); A.D.M. was supported by MRC (MR/T030062/1); A.M.M was supported by the University of Bristol; K.D.A. was supported by the Catto Charitable Trust; Sensititre plates were provided by Thermo Fisher Scientific Ltd.; we thank Duncan Porter and Toby Hampshire for their helpful discussions and for assisting with the selection and sourcing of the plates. M.E.K.H. as an Academic Clinical Lecturer was funded by the NIHR. The views expressed in this publication are those of the author(s) and not necessarily those of the NIHR, NHS, or the UK Department of Health and Social Care.

## Data availability

All sequence data described herein have been submitted to the European Nucleotide Archive (http://www.ebi.ac.uk/ena) and are available under the accession numbers PRJEB92194 and PRJEB97996. Individual accessions are provided in the supplementary tables. Code is available from TBD.

## Author Contributions

M.R.J.C., A.D.M., M.E.KH., and K.D.A. conceived the study. K.D.A. performed the experiments. E.J., N.B., M.Akter, M.Attwood., J.M., and J.M.S. provided phages and bacterial isolates. M.T. developed the ImageJ script for quantitative image analysis. K.D.A, S.M., and C.W. analysed the data. K.D.A. with O.G., T.S.P., A.D.M., and M.R.J.C drafted the manuscript. M.R.J.C., A.D.M., T.S.P., and M.E.K.H. supervised the work. Data were generated by K.D.A, S.M., C.W., and A.D.

All authors have read and approved the manuscript.

## References

1. Antimicrobial Resistance Collaborators. Global burden of bacterial antimicrobial resistance in 2019: a systematic analysis. Lancet 399, 629–655 (2022).

2. Kucers’ the Use of Antibiotics : A Clinical Review of Antibacterial, Antifungal, Antiparasitic, and Antiviral Drugs. (CRC Press, Philadelphia, PA).

3. Naber, K. G., Tirán-Saucedo, J., Wagenlehner, F. M. E. & RECAP group. Psychosocial burden of recurrent uncomplicated urinary tract infections. GMS Infect. Dis. 10, Doc01 (2022).

4. Gould, C. V. et al. Guideline for prevention of catheter-associated urinary tract infections 2009. Infect. Control Hosp. Epidemiol. 31, 319–326 (2010).

5. WHO bacterial priority pathogens list, 2024: Bacterial pathogens of public health importance to guide research, development and strategies to prevent and control antimicrobial resistance. https://www.who.int/publications/i/item/9789240093461 (2024).

6. Antimicrobial resistance surveillance in Europe 2022 - 2020 data. European Centre for Disease Prevention and Control https://www.ecdc.europa.eu/en/publications-data/antimicrobial-resistance-surveillance-europe-2022-2020-data (2022).

7. Adams-Sapper, S., Diep, B. A., Perdreau-Remington, F. & Riley, L. W. Clonal composition and community clustering of drug-susceptible and -resistant Escherichia coli isolates from bloodstream infections. Antimicrob. Agents Chemother. 57, 490–497 (2013).

8. Choi, H. J. et al. Characteristics of Escherichia coli urine isolates and risk factors for secondary bloodstream infections in patients with urinary tract infections. Microbiol. Spectr. 10, e0166022 (2022).

9. Ciesielczuk, H. et al. Trends in ExPEC serogroups in the UK and their significance. Eur. J. Clin. Microbiol. Infect. Dis. 35, 1661–1666 (2016).

10. Day, M. J. et al. Extended-spectrum β-lactamase-producing Escherichia coli in human-derived and foodchain-derived samples from England, Wales, and Scotland: an epidemiological surveillance and typing study. Lancet Infect. Dis. 19, 1325–1335 (2019).

11. Dale, A. P. et al. Genomes of Escherichia coli bacteraemia isolates originating from urinary tract foci contain more virulence-associated genes than those from non-urinary foci and neutropaenic hosts. J. Infect. 77, 534–543 (2018).

12. Gibreel, T. M. et al. Population structure, virulence potential and antibiotic susceptibility of uropathogenic Escherichia coli from Northwest England. J. Antimicrob. Chemother. 67, 346–356 (2012).

13. Holmbom, M. et al. Risk factors and outcome due to extended-spectrum β-lactamase-producing uropathogenic Escherichia coli in community-onset bloodstream infections: A ten-year cohort study in Sweden. PLoS One 17, e0277054 (2022).

14. Horner, C. et al. Escherichia coli bacteraemia: 2 years of prospective regional surveillance (2010-12). J. Antimicrob. Chemother. 69, 91–100 (2014).

15. Kallonen, T. et al. Systematic longitudinal survey of invasive Escherichia coli in England demonstrates a stable population structure only transiently disturbed by the emergence of ST131. Genome Res. (2017) doi:10.1101/gr.216606.116.

16. Li, D. et al. Dominance of Escherichia coli sequence types ST73, ST95, ST127 and ST131 in Australian urine isolates: a genomic analysis of antimicrobial resistance and virulence linked to F plasmids. Microb. Genom. 9, (2023).

17. Lau, S. H. et al. Major uropathogenic Escherichia coli strain isolated in the northwest of England identified by multilocus sequence typing. J. Clin. Microbiol. 46, 1076–1080 (2008).

18. Lipworth, S. et al. Ten-year longitudinal molecular epidemiology study of Escherichia coli and Klebsiella species bloodstream infections in Oxfordshire, UK. Genome Med. 13, 144 (2021).

19. Loc-Carrillo, C. & Abedon, S. T. Pros and cons of phage therapy. Bacteriophage 1, 111–114 (2011).

20. Ryan, E. M., Alkawareek, M. Y., Donnelly, R. F. & Gilmore, B. F. Synergistic phage-antibiotic combinations for the control of Escherichia coli biofilms in vitro. FEMS Immunol. Med. Microbiol. 65, 395–398 (2012).

21. Shlezinger, M., Coppenhagen-Glazer, S., Gelman, D., Beyth, N. & Hazan, R. Eradication of vancomycin-resistant enterococci by combining phage and vancomycin. Viruses 11, 954 (2019).

22. Grygorcewicz, B. et al. Environmental phage-based cocktail and antibiotic combination effects on Acinetobacter baumannii biofilm in a human urine model. Microb. Drug Resist. 27, 25–35 (2021).

23. Xu, W. et al. The identification of phage vB_1086 of multidrug-resistant Klebsiella pneumoniae and its synergistic effects with ceftriaxone. Microb. Pathog. 171, 105722 (2022).

24. Kebriaei, R. et al. Optimization of Phage-Antibiotic Combinations against Staphylococcus aureus Biofilms. Microbiol. Spectr. 11, e0491822 (2023).

25. Wilburn, K., Matrishin, C. B., Choudhury, A., Larsen, R. & Wildschutte, H. Tradeoffs between evolved phage resistance and antibiotic susceptibility in a highly drug-resistant cystic fibrosis-derived Pseudomonas aeruginosa strain. Phage (New Rochelle) 5, 45–52 (2024).

26. Simon, K., Pier, W., Krüttgen, A. & Horz, H.-P. Synergy between phage Sb-1 and oxacillin against methicillin-resistant Staphylococcus aureus. Antibiotics (Basel) 10, 849 (2021).

27. Zuo, P., Yu, P. & Alvarez, P. J. J. Aminoglycosides antagonize bacteriophage proliferation, attenuating phage suppression of bacterial growth, biofilm formation, and antibiotic resistance. Appl. Environ. Microbiol. 87, e0046821 (2021).

28. Luo, C.-H., Hsu, Y.-H., Wu, W.-J., Chang, K.-C. & Yeh, C.-S. Phage digestion of a bacterial capsule imparts resistance to two antibiotic agents. Microorganisms 9, 794 (2021).

29. Nang, S. C. et al. Phage resistance in Klebsiella pneumoniae and bidirectional effects impacting antibiotic susceptibility. Clin. Microbiol. Infect. 30, 787–794 (2024).

30. Ribes-Martínez, L., Muñoz-Egea, M.-C., Yuste, J., Esteban, J. & García-Quintanilla, M. Bacteriophage therapy as a promising alternative for antibiotic-resistant Enterococcus faecium: Advances and challenges. Antibiotics (Basel) 13, (2024).

31. Forde, B. M. et al. Population dynamics of an Escherichia coli ST131 lineage during recurrent urinary tract infection. Nat. Commun. 10, 3643 (2019).

32. Criscuolo, A. A fast alignment-free bioinformatics procedure to infer accurate distance-based phylogenetic trees from genome assemblies. Res. Ideas Outcomes 5, (2019).

33. Nishimura, Y. et al. ViPTree: The viral proteomic tree server. Bioinformatics (2017) doi:10.1093/bioinformatics/btx157.

34. Picard, B. et al. The link between phylogeny and virulence in Escherichia coli extraintestinal infection. Infect. Immun. 67, 546–553 (1999).

35. Gaborieau, B. et al. Prediction of strain level phage-host interactions across the Escherichia genus using only genomic information. Nat. Microbiol. 9, 2847–2861 (2024).

36. Rodrigues, C. et al. Description of Klebsiella africanensis sp. nov., Klebsiella variicola subsp. tropicalensis subsp. nov. and Klebsiella variicola subsp. variicola subsp. nov. Res. Microbiol. 170, 165–170 (2019).

37. Andersson, D. I., Nicoloff, H. & Hjort, K. Mechanisms and clinical relevance of bacterial heteroresistance. Nat. Rev. Microbiol. 17, 479–496 (2019).

38. Clinical Breakpoint Tables. The European Committee on Antimicrobial Susceptibility Testing - EUCAST https://www.eucast.org/bacteria/clinical-breakpoints-and-interpretation/clinical-breakpoint-tables/.

39. Gonzalez, M. A. et al. Multiple-dose pharmacokinetics and safety of ciprofloxacin in normal volunteers. Antimicrob. Agents Chemother. 26, 741–744 (1984).

40. Novelli, A. & Rosi, E. Pharmacological properties of oral antibiotics for the treatment of uncomplicated urinary tract infections. J. Chemother. 29, 10–18 (2017).

41. Kim, M. et al. Phage-antibiotic synergy via delayed lysis. Appl. Environ. Microbiol. 84, (2018).

42. Moradpour, Z., Yousefi, N., Sadeghi, D. & Ghasemian, A. Synergistic bactericidal activity of a naturally isolated phage and ampicillin against urinary tract infecting Escherichia coli O157. Iran. J. Basic Med. Sci. 23, 257–263 (2020).

43. Easwaran, M., De Zoysa, M. & Shin, H.-J. Application of phage therapy: Synergistic effect of phage EcSw (ΦEcSw) and antibiotic combination towards antibiotic-resistant Escherichia coli. Transbound. Emerg. Dis. 67, 2809–2817 (2020).

44. Gu Liu, C., et al. Phage-antibiotic synergy is driven by a unique combination of antibacterial mechanism of action and stoichiometry. MBio 11, (2020).

45. Bulssico, J., PapukashvilI, I., Espinosa, L., Gandon, S. & Ansaldi, M. Phage-antibiotic synergy: Cell filamentation is a key driver of successful phage predation. PLoS Pathog. 19, e1011602 (2023).

46. Pacios, O. et al. Enhanced antibacterial activity of repurposed mitomycin C and imipenem in combination with the lytic phage vB_KpnM-VAC13 against clinical isolates of Klebsiella pneumoniae. Antimicrob. Agents Chemother. 65, e0090021 (2021).

47. Michodigni, N. F. et al. Formulation of phage cocktails and evaluation of their interaction with antibiotics in inhibiting carbapenemase-producing Klebsiella pneumoniae in vitro in Kenya. Afr. J. Lab. Med. 11, 1803 (2022).

48. Ziller, L. et al. Newly isolated Drexlerviridae phage LAPAZ is physically robust and fosters eradication of Klebsiella pneumoniae in combination with meropenem. Virus Res. 347, 199417 (2024).

49. Qin, K. et al. Phage-antibiotic synergy suppresses resistance emergence of Klebsiella pneumoniae by altering the evolutionary fitness. MBio 15, e0139324 (2024).

50. Chan, B. K. et al. Phage selection restores antibiotic sensitivity in MDR Pseudomonas aeruginosa. Sci. Rep. 6, 26717 (2016).

51. Koderi Valappil, S., et al. Survival comes at a cost: A coevolution of phage and its host leads to phage resistance and antibiotic sensitivity of Pseudomonas aeruginosa multidrug resistant strains. Front. Microbiol. 12, 783722 (2021).

52. Köhler, T., Michea-Hamzehpour, M., Epp, S. F. & Pechere, J. C. Carbapenem activities against Pseudomonas aeruginosa: respective contributions of OprD and efflux systems. Antimicrob. Agents Chemother. 43, 424–427 (1999).

53. Kever, L. et al. Aminoglycoside antibiotics inhibit phage infection by blocking an early step of the infection cycle. MBio 13, e0078322 (2022).

54. Verma, V., Harjai, K. & Chhibber, S. Structural changes induced by a lytic bacteriophage make ciprofloxacin effective against older biofilm of Klebsiella pneumoniae. Biofouling 26, 729–737 (2010).

55. Jung, Y., Kim, J. & Ryu, S. Antagonistic effect of kanamycin on kanamycin-resistant Escherichia coli infection by BtuB-targeting bacteriophages. Arch. Virol. 170, 172 (2025).

56. Vukovic, D. et al. Antibacterial potential of non-tailed icosahedral phages alone and in combination with antibiotics. Curr. Microbiol. 81, 215 (2024).

57. Wang, Z. et al. Combination therapy of phage vB_KpnM_P-KP2 and gentamicin combats acute pneumonia caused by K47 serotype Klebsiella pneumoniae. Front. Microbiol. 12, 674068 (2021).

58. Paranos, P. et al. Reversal of phenotypic resistance in multi-drug resistant carbapenemase-producing K. pneumoniae clinical isolates due to in vitro synergistic interactions between bacteriophages and antibiotics at clinically achievable concentrations. J. Antimicrob. Chemother. 80, 1997–2006 (2025).

59. Sanchez, B. C. et al. Development of phage cocktails to treat E. coli catheter-associated urinary tract infection and associated biofilms. Front. Microbiol. 13, 796132 (2022).

60. Shein, A. M. S. et al. Phage cocktail amikacin combination as a potential therapy for bacteremia associated with carbapenemase producing colistin resistant Klebsiella pneumoniae. Sci. Rep. 14, 28992 (2024).

61. Vera-Mansilla, J., Silva-Valenzuela, C. A., Sánchez, P. & Molina-Quiroz, R. C. Bacteriophages potentiate the effect of antibiotics by eradication of persister cells and killing of biofilm-forming cells. Res. Microbiol. 174, 104083 (2023).

62. Wang, L., Tkhilaishvili, T., Bernal Andres, B., Trampuz, A. & Gonzalez Moreno, M. Bacteriophage-antibiotic combinations against ciprofloxacin/ceftriaxone-resistant Escherichia coli in vitro and in an experimental Galleria mellonella model. Int. J. Antimicrob. Agents 56, 106200 (2020).

63. Khunti, P. et al. A novel coli myophage and antibiotics synergistically inhibit the growth of the uropathogenic E. coli strain CFT073 in stoichiometric niches. Microbiol. Spectr. e0088923 (2023).

64. Lopes, A., Pereira, C. & Almeida, A. Sequential combined effect of phages and antibiotics on the inactivation of Escherichia coli. Microorganisms 6, 125 (2018).

65. Gorodnichev, R. B. et al. Phage-antibiotic combinations against Klebsiella pneumoniae: impact of methodological approaches on effect evaluation. Front. Microbiol. 16, 1530819 (2025).

66. Malik, S., Nehra, K., Mann, A., Jagdish, R. & Rana, J. S. Characterization and synergy studies of Caudoviricete Escherichia phage FS2B infecting multi-drug resistant uropathogenic Escherichia coli isolates. Int. Microbiol. 27, 155–166 (2024).

67. Morris, T. C., Reyneke, B., Khan, S. & Khan, W. Phage-antibiotic synergy to combat multidrug resistant strains of Gram-negative ESKAPE pathogens. Sci. Rep. 15, 17235 (2025).

68. Van Twest, R. & Kropinski, A. M. Bacteriophage enrichment from water and soil. Methods Mol. Biol. 501, 15–21 (2009).

69. Bacteriophages: Biology, Technology, Therapy. (Springer International Publishing, Cham, Switzerland, 2021).

70. Alcock, B. P. et al. CARD 2023: expanded curation, support for machine learning, and resistome prediction at the Comprehensive Antibiotic Resistance Database. Nucleic Acids Res. 51, D690–D699 (2023).

71. Magiorakos, A.-P. et al. Multidrug-resistant, extensively drug-resistant and pandrug-resistant bacteria: an international expert proposal for interim standard definitions for acquired resistance. Clin. Microbiol. Infect. 18, 268–281 (2012).

72. Mazzocco, A., Waddell, T. E., Lingohr, E. & Johnson, R. P. Enumeration of bacteriophages by the direct plating plaque assay. Methods Mol. Biol. 501, 77–80 (2009).

73. ThermoScientific. 2019. Instructions for Use: Thermo Scientific Sensititre MIC Susceptibility Plates for Testing Non-Fastidious Gram Negative and Gram Positive Isolates, 037-NFAST FDA-USA Only. *CID10253 Ed*.

74. Schindelin, J., et al. Fiji: an open-source platform for biological-image analysis. Nat. Methods 9, 676–682 (2012).

75. Benjamini, Y., Krieger, A. M. & Yekutieli, D. Adaptive linear step-up procedures that control the false discovery rate. Biometrika 93, 491–507 (2006).

76. Rihtman, B., Meaden, S., Clokie, M. R. J., Koskella, B. & Millard, A. D. Assessing Illumina technology for the high-throughput sequencing of bacteriophage genomes. PeerJ 4, e2055 (2016).

77. Michniewski, S., et al. Rolling out *plaque-2-sequence* : a single plaque sequencing approach enabling rapid, low-cost sequencing of phages directly from plaques. bioRxiv (2025) doi:10.1101/2025.11.01.684647.

78. Jolley, K. A., Bray, J. E. & Maiden, M. C. J. Open-access bacterial population genomics: BIGSdb software, the PubMLST.org website and their applications. Wellcome Open Res. 3, 124 (2018).

79. Alikhan, N. F., Zhou, Z., Sergeant, M. J. & Achtman, M. A genomic overview of the population structure of Salmonella. PLoS Genetics Preprint at 10.1371/journal.pgen.1007261 (2018).

80. Waters, N. R., Abram, F., Brennan, F., Holmes, A. & Pritchard, L. Easy phylotyping of Escherichia coli via the EzClermont web app and command-line tool. Access Microbiol. 2, acmi000143 (2020).

81. Clermont, O., Christenson, J. K., Denamur, E. & Gordon, D. M. The Clermont Escherichia coli phylo-typing method revisited: improvement of specificity and detection of new phylo-groups. Environ. Microbiol. Rep. 5, 58–65 (2013).

82. Hennart, M. et al. A dual barcoding approach to bacterial strain nomenclature: Genomic taxonomy of Klebsiella pneumoniae strains. Mol. Biol. Evol. 39, (2022).

83. Ondov, B. D. et al. Mash: fast genome and metagenome distance estimation using MinHash. Genome Biol. 17, 132 (2016).

84. Guénoche, A. & Garreta, H. Can we have confidence in a tree representation? in Computational Biology 45–56 (Springer Berlin Heidelberg, Berlin, Heidelberg, 2001).

85. Zhou, Z. et al. GrapeTree: visualization of core genomic relationships among 100,000 bacterial pathogens. Genome Res. 28, 1395–1404 (2018).

86. Yu, G., Lam, T. T.-Y., Zhu, H. & Guan, Y. Two methods for mapping and visualizing associated data on phylogeny using ggtree. Mol. Biol. Evol. 35, 3041–3043 (2018).

87. Xu, S. et al. GgtreeExtra: Compact visualization of richly annotated phylogenetic data. Mol. Biol. Evol. 38, 4039–4042 (2021).

88. Galardini, M. et al. Phenotype inference in an Escherichia coli strain panel. Elife 6, (2017).

89. Shen, A. & Millard, A. Phage genome annotation: Where to begin and end. PHAGE 2, 183–193 (2021).

90. Bankevich, A. et al. SPAdes: A New Genome Assembly Algorithm and Its Applications to Single-Cell Sequencing. J. Comput. Biol. 19, 455–477 (2012).

91. Prjibelski, A., Antipov, D., Meleshko, D., Lapidus, A. & Korobeynikov, A. Using SPAdes DE Novo Assembler. Curr. Protoc. Bioinformatics 70, e102 (2020).

92. Seemann, T. Prokka: rapid prokaryotic genome annotation. Bioinformatics 30, 2068–2069 (2014).

93. Terzian, P., et al. PHROG: families of prokaryotic virus proteins clustered using remote homology. NAR Genom Bioinform 3, lqab067 (2021).

94. Millard, A. et al. TaxMyPhage: Automated taxonomy of dsDNA phage genomes at the genus and species level. Phage (New Rochelle) (2025) doi:10.1089/phage.2024.0050.

95. Marçais, G. et al. MUMmer4: A fast and versatile genome alignment system. PLoS Comput. Biol. 14, e1005944 (2018).

96. Page, A. J. et al. Roary: rapid large-scale prokaryote pan genome analysis. Bioinformatics 31, 3691–3693 (2015).

97. Madeira, F. et al. The EMBL-EBI Job Dispatcher sequence analysis tools framework in 2024. Nucleic Acids Res. 52, W521–W525 (2024).

98. Turner, D. et al. Summary of taxonomy changes ratified by the International Committee on Taxonomy of Viruses (ICTV) from the Bacterial Viruses Subcommittee, 2025. J. Gen. Virol. 106, 002111 (2025).

