## Supplemental Figures for "Large scale phage-antibiotic combination studies reveal key combinations for urinary tract infection and urosepsis treatments"

### Supplemental Figure 1 – Phylogenetic Trees and ANI Matrix, *Tequatroviruses*

Phylogenetic tree and similarity matrix for the eight phages of the *Tequatrovirus* genus included in this study, with phages 112 and 130 (>99.9% ANI genomic similarity) highlighted.

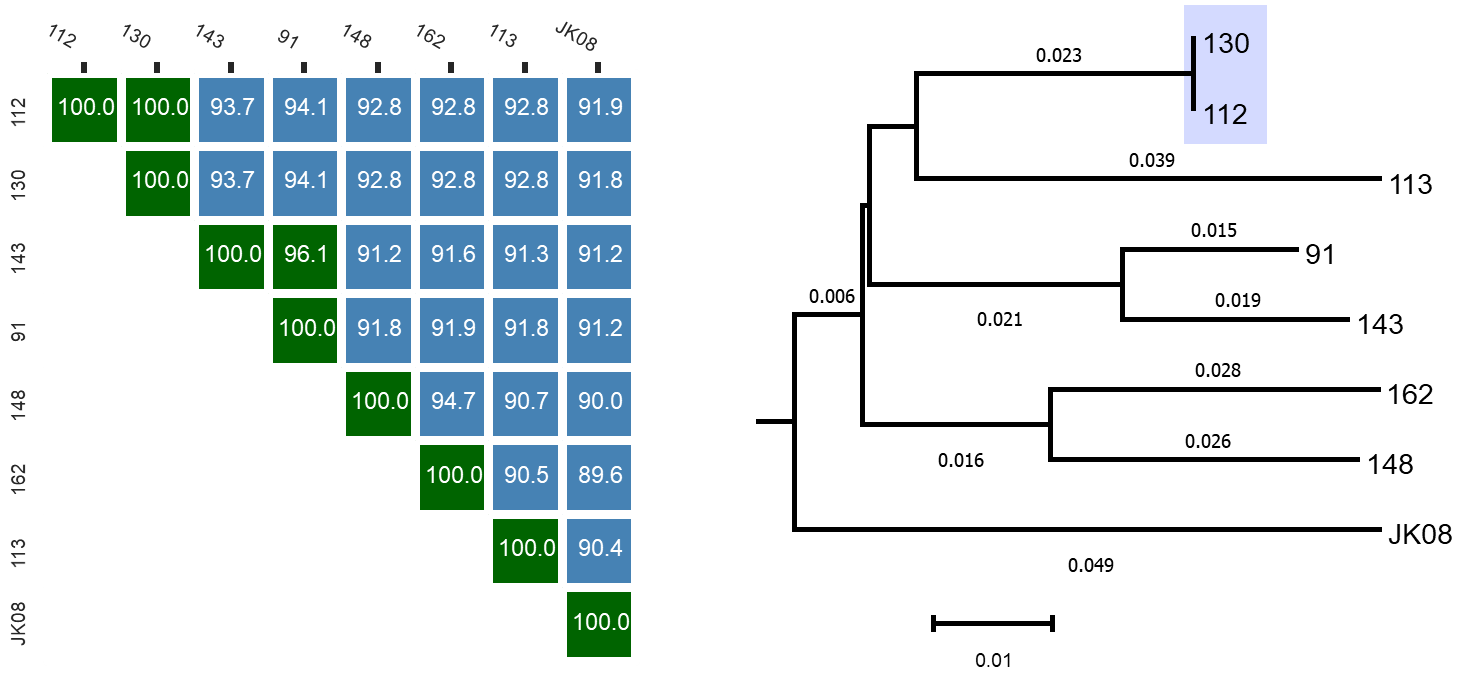

### Supplemental Figure 2 – Phylogenetic Trees and ANI Matrix, *Felixounaviruses*

Phylogenetic tree and similarity matrix for the three phages of the *Felixounavirus* genus included in this study, with phages JK19 and JK21 (>99.9% ANI genomic similarity) highlighted.

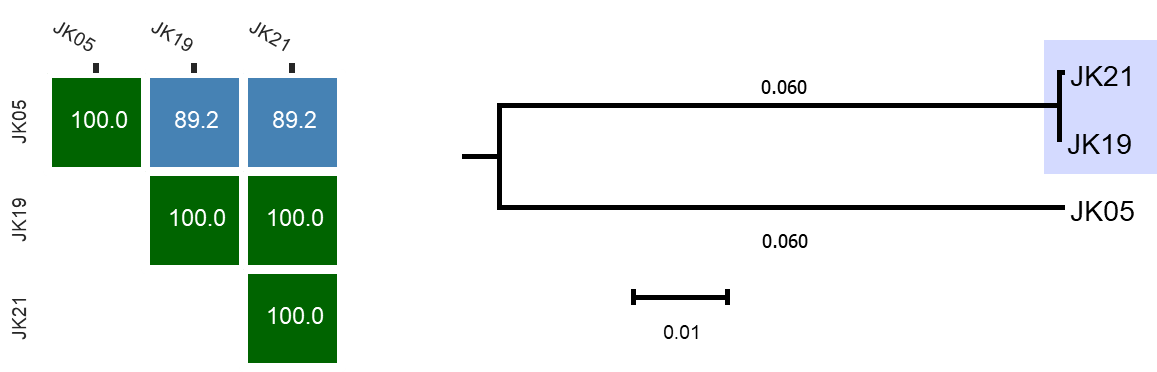

### Supplemental Figure 3 – Phylogenetic Trees and ANI Matrix, *Sugarlandviruses*

Phylogenetic tree and similarity matrix for the eight phages of the *Sugarlandvirus* genus included in this study, with phages BMCPR_006077, Ponyo_var3, BS1_var2, PG2, RedSea1_var2, and EJMex1 (>99.5% ANI genomic similarity) highlighted.

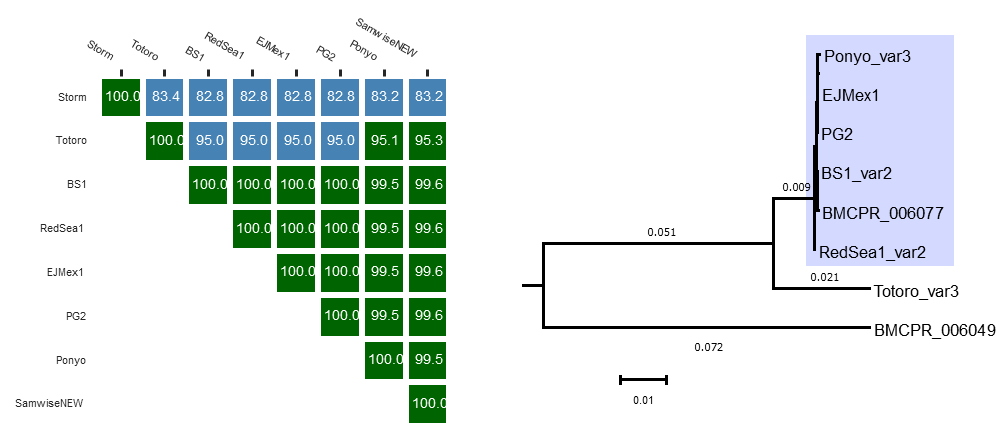

### Supplemental Figure 4 – Phylogenetic Trees and ANI Matrix, *Eowynviruses*

Phylogenetic tree and similarity matrix for the three phages of the *Eowynvirus* genus included in this study, with phages Eowyn_var3 and Spernal1_var3 (>99.9% ANI genomic similarity) highlighted.

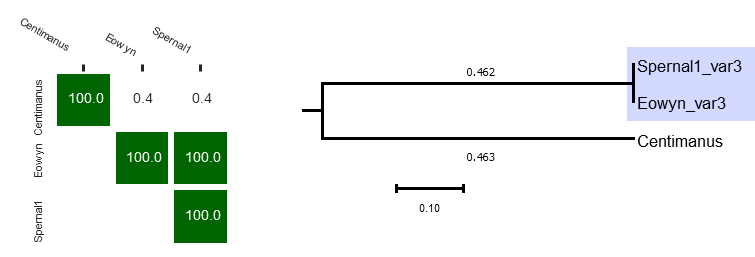

### Supplemental Figure 5 – Phylogenetic Trees and ANI Matrix, *Taipeiviruses*

Phylogenetic tree and similarity matrix for the three phages of the *Taipeivirus* genus included in this study, with phages LilBean_var3 and Bilbo_var3 (>99.9% ANI genomic similarity) highlighted.

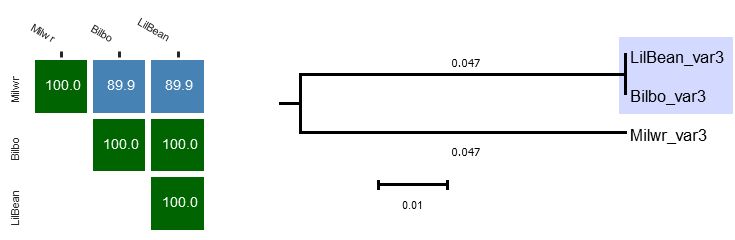

### Supplemental Figure 6 – Scoring Schematic for Screening Assays

Scoring schematic for the broth-based screening assay (left) and agar-based spot test assay (right). Representative examples illustrating the range of scores for each assay are shown.

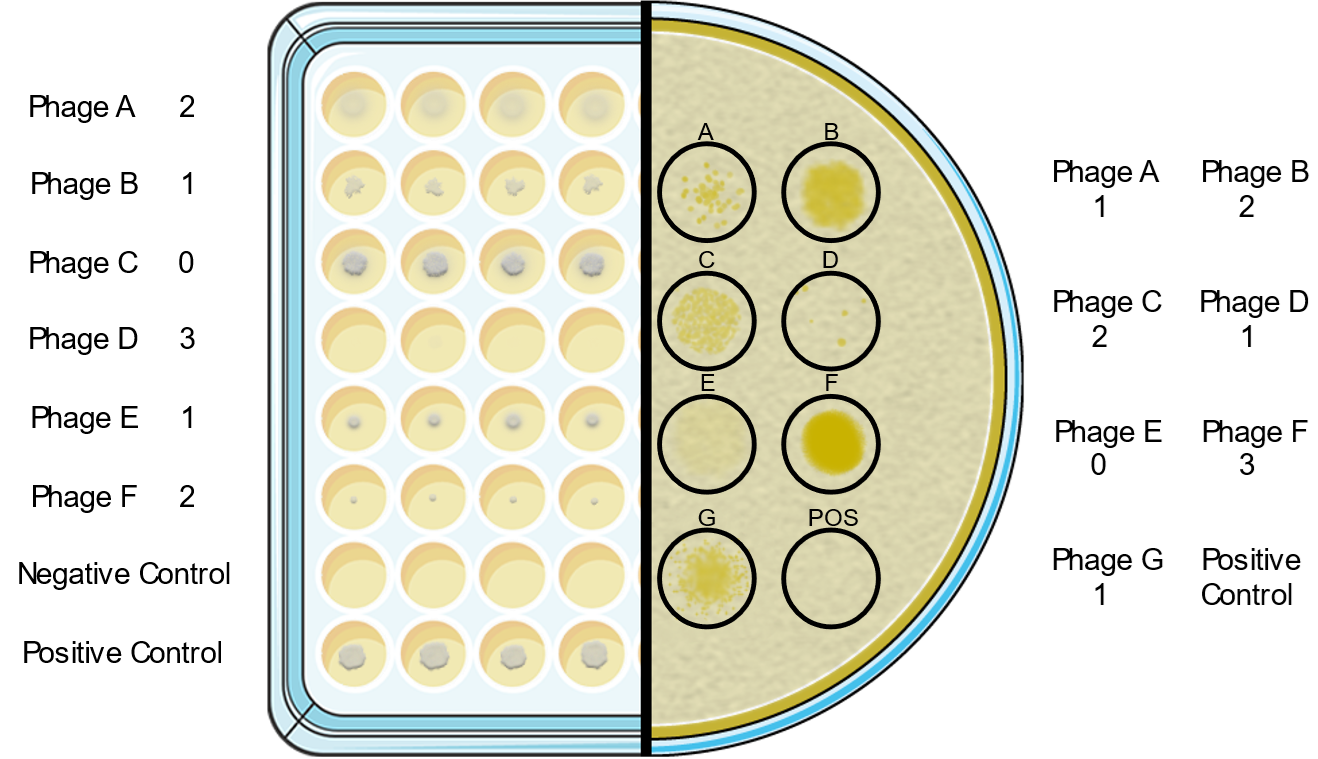
